# Critical slowing as a biomarker for seizure susceptibility

**DOI:** 10.1101/689893

**Authors:** Matias I. Maturana, Christian Meisel, Katrina Dell, Philippa J. Karoly, Wendyl D’Souza, David B. Grayden, Anthony N. Burkitt, Premysl Jiruska, Jan Kudlacek, Jaroslav Hlinka, Mark J. Cook, Levin Kuhlmann, Dean R. Freestone

## Abstract

The human brain has the capacity to rapidly change state, and in epilepsy these state changes can be catastrophic, resulting in loss of consciousness, injury and even death. Theoretical interpretations considering the brain as a dynamical system would suggest that prior to a seizure recorded brain signals may exhibit critical slowing, a warning signal preceding many critical transitions in dynamical systems. Using long-term intracranial electroencephalography (iEEG) recordings from fourteen patients with focal epilepsy, we found key signatures of critical slowing prior to seizures. Signals related to a critically slowing process fluctuated over temporally long scales (hours to days), longer than would be detectable in standard clinical evaluation settings. Seizure risk was associated with a combination of these signals together with epileptiform discharges. These results provide strong validation of theoretical models and demonstrate that critical slowing is a reliable indicator that could be used in seizure forecasting algorithms.

## Introduction

The unexpected nature of epileptic seizures represents the major clinical disability of epilepsy [1]. The mechanisms underlying the transition from a normal to a seizure state are currently an open question [2–4]. Unravelling the mechanisms underlying seizure generation could form the basis of much needed new treatment strategies, particularly for patients where existing treatments are ineffective.

Abrupt state changes in natural systems, including the onset of seizures, can be due to critical transitions [5]. A characteristic of a system that is approaching a critical transition is a phenomenon called “critical slowing”, which refers to the tendency of a system to take longer to return to equilibrium after perturbations, indicated by an increase in signal variance and autocorrelation. Critical slowing has been observed in many systems, including cell population collapse in bacterial cultures [6], crashes in financial markets [7], and earthquakes [8]. Critical transitions have been employed to describe neural systems, such as onset of depression [9], changes in perception [10], switching between motor programs [11], onset of spiking in neurons [12], and termination of epileptic seizures [13].

It has been hypothesized that a rapid transition from normal brain activity to an epileptic seizure is such a critical transition [3, 4, 14–17]. However, there is insufficient empirical evidence to test this hypothesis in humans, because of the lack of long-term recordings as well as intra-patient and inter-patient variability. Empirical validation of critical slowing in humans would provide vital support for current theoretical models of seizure generation and of the dynamics of the brain in general. Furthermore, it could aid in forecasting seizures and potential titration of epilepsy therapies.

Computational neural models are powerful tools for studying the dynamics of the brain. Numerous computational models of epilepsy suggest that critical slowing occurs prior to seizures [14, 16, 18]. Mathematical analyses of dynamic systems, combined with simulations, enable classification of bifurcations and critical transitions. Simulations enable controlled experiments that vary the parameters of the model and reveal statistical markers that are representative of transition susceptibility, such as increases in signal variance and autocorrelation [13]. While some methods have been developed to track control parameters from clinically captured EEG in epilepsy [19, 20], this approach is not straightforward. Alternatively, tracking the statistical markers in clinical EEG recordings may constitute a direct test of the hypothesis that seizures are critical transitions.

In this paper we test the hypothesis that critical slowing is a biomarker of seizure susceptibility. We examine hallmark signals of critical slowing using a continuous iEEG dataset from the first-in-human trial of an implanted seizure prediction device that was recorded over multiple years [21]. As critical slowing can potentially manifest at very long timescales, the long duration dataset used for this analysis provides a unique opportunity where critical slowing in humans can be robustly investigated. We show that the autocorrelation and variance of the iEEG signals are modulated by patient-specific cycles over long temporal scales. Furthermore, we show that modulations of the variance and autocorrelation are related to seizure susceptibility – a probabilistic propensity to have seizures – supporting the hypothesis that seizures constitute a critical transition.

## Results

### Conceptualization of critical slowing in epilepsy

In epilepsy, seizure events could be described as a “phase transition” or “critical transition”, based on deterministic or mechanistic dynamics, where the brain shifts from a normal to a seizure state [5]. Theoretical predictions from dynamical systems models suggest that critical transitions are preceded by increases in signal variance and autocorrelation prior to state changes. There are numerous models that can describe state changes and there are many possible paths leading to a state change [16]. It may be possible to detect the occurrence of critical slowing prior to a seizure [3], and we describe here a model to demonstrate how critical slowing may occur with the view that this concept may hold for a wider class of models.

Figure 1 illustrates our conceptualization of critical slowing in the context of iEEG signals. The iEEG signal is thought to primarily originate from the population activity of pyramidal cells [22, 23], whose activity is modulated by interactions with pre-synaptic neurons. One way to conceptualize critical slowing relative to the iEEG signal is by considering the dynamics of the signal. The mean action potential firing rate of a population of pyramidal cells is modulated by the cellular integration of post-synaptic potentials. Typical parameters determining the mean output firing rate are the amplitudes and time constants of the post synaptic potentials, the nonlinear threshold for firing, and the slope of the nonlinearity [19]. On a fast time scale, these parameters can be considered constant. However, on a much slower temporal scale, these parameters may vary due to effects of neuromodulators and hormonal and other biological processes, which in turn affect the behavior of the neural population [24].

**Figure 1:**
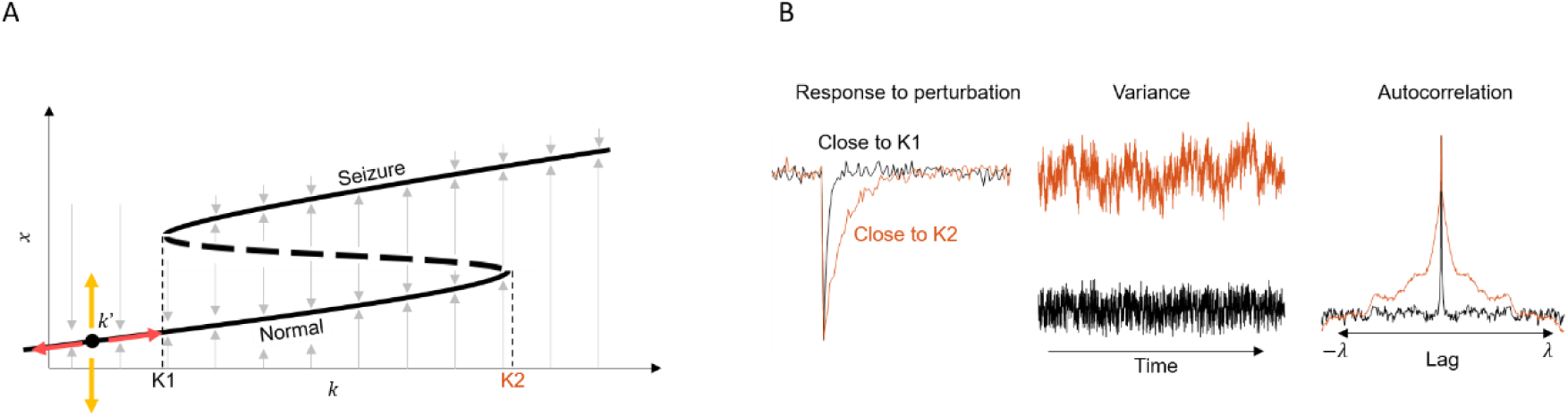
Conceptualization of critical slowing with regards to intracranial EEG (iEEG) signals. A) A bifurcation diagram to show how *fast* perturbations in *x* can cause a transition into the seizure state. The state *x* represents a property of the iEEG signal, which could be the mean action potential firing rate or mean membrane potential of pyramidal cells (believed to be the principle generators of the iEEG signal). A differential equation governs the steady-state solutions of *x* for any *slow* parameter *k*. The steady-state solutions are represented by the thick black lines. As *k* is varied in the positive direction, *x* transitions from having one (normal brain activity) steady-state solution, to three solutions between K1 and K2: one normal, one unstable (dashed line), then to one seizure state. B) Different signal dynamics are observed near points K1 (black) and K2 (orange). Close to K2, the system time constant is greater, resulting in an increased recovery time (i.e. slowing) to perturbations. This results in an increase in signal variance and an autocorrelation function with a longer time constant. Critical slowing refers to the idea that signal slowing occurs when the system parameter *k* is close to a critical transition or bifurcation point (i.e. K1 and K2).

Several models of epilepsy describe the change in brain state from normal to seizure as a bifurcation that occurs as the system crosses a critical point [14–17, 25, 26]. Figure 1A illustrates a one-dimensional dynamical system, where the state *x* is modulated by parameter *k*. We can think of *x* as being a fast-changing property of the iEEG signal; for example, it could represent the time-varying mean membrane potential of pyramidal cells averaged locally in space. The parameter *k* represents the slow driving element, which could represent the response of the brain to a variety of factors such as medication, sleep, or metabolic processes. The thick black lines in Figure 1A represent the steady-state or equilibrium values taken by *x* for any given value of *k*.

Values of *k* between K1 and K2 represent solutions to the dynamical system where *x* can take on three steady states. The unstable state (dashed line) is a state from which the system quickly diverges to one of the two stable states – the *normal* and *seizure* states. As *k* approaches K2, the unstable state and normal state converge, as the dominant eigenvalue solutions converge at zero. Simply put, the system’s rate of change near the critical point approaches zero, and becomes increasingly slower in recovering from perturbations. One way to measure this effect is to track the signal’s short-term variance and autocorrelation (a measure of a signal’s similarity to its past).

Figure 1B illustrates the observations expected with critical slowing. For values of *k* near K1, the probability of a transition from the normal state to the seizure state is low, as the energy required to cross the unstable state into the seizure state is large, and the system rapidly returns to equilibrium after a perturbation. However, as *k* approaches K2, the energy of a perturbation required to cross the unstable boundary is reduced, resulting in a higher probability of a state transition, and longer time to recover from a perturbation.

In this work, we use the markers of critical slowing as a proxy to track the slow parameter *k* that, in turn, offers a biomarker for seizure susceptibility [3]. Moreover, we focus on seizure forecasting [3, 27] rather than seizure prediction [2], as we do not attempt to predict the fast (noise) event that perturbs the system towards the seizure state.

### Data summary

Continuous iEEG from 15 patients was used in this study [21]. For each patient, an array of 16 electrodes was placed on the surface of the brain near the presumed epileptogenic zone (e.g. Figure 2A). After pre-processing (see Methods), a total of 2871 seizures were analyzed (see Table 1), with gaps in the data comprising 26% of the total data (minimum: 1.25%, maximum: 40.11%). Only clinically correlated and clinically equivalent seizures, as defined by Cook et al. [21], were considered.

**Figure 2:**
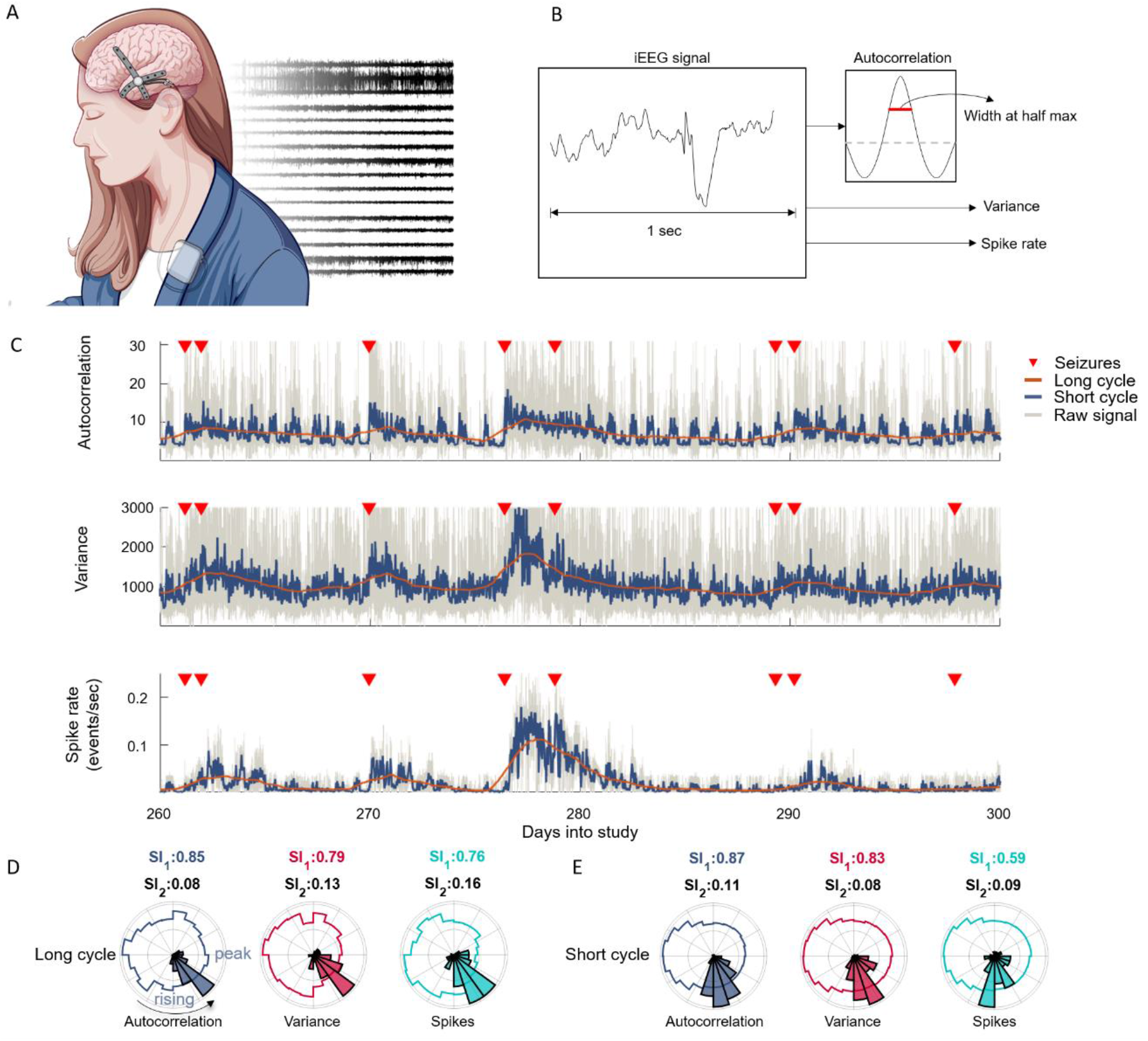
Details of analysis of Patient 1. A) An illustration of the implanted electrodes that captured continuous EEG signals from the surface of the brain at 16 different locations, with examples of the signals. B) Autocorrelation, variance, and spike rates were computed from one second segments of iEEG every two minutes. The autocorrelation was taken as the width at half maximum of the autocorrelation function. C) The autocorrelation (top), variance (middle) and spike rate signals were filtered using a moving average filter to reveal short and long rhythms. Seizures (red triangles) preferentially occurred on the rising phases of the signals. D) Normalized polar histograms of the seizure phases for the long cycles demonstrated that most seizures occurred on a narrow phase of the autocorrelation (grey), variance (red) and spike rate (cyan) signals. In each plot the rising phase and signal peak is as denoted in the autocorrelation polar plot. The synchronization indices for the signals (black) and histograms (colored) are shown above each plot. E) Normalized polar histograms of the seizure phases for the short cycles.

**Table 1:** Patient data summary. The total number of seizures, mean seizure rate, the number of recorded days and the percentage of data dropouts are tabulated.

|  |  |  |  |  |
| --- | --- | --- | --- | --- |
| Patient | Total seizures | Mean seizure rate (seizures/day) | Number of days | Data dropout (%) |
| 1 | 151 | 0.20 | 767 | 35.1 |
| 2 | 32 | 0.04 | 730 | 19.6 |
| 3* | 374 | 0.70 | 557 | 78.3 |
| 4 | 22 | 0.09 | 233 | 26.2 |
| 5 | 9 | 0.03 | 273 | 40.11 |
| 6 | 71 | 0.16 | 441 | 2.95 |
| 7 | 313 | 1.70 | 185 | 11.93 |
| 8 | 466 | 0.84 | 558 | 35.7 |
| 9 | 202 | 0.51 | 395 | 36.4 |
| 10 | 545 | 1.46 | 373 | 6.9 |
| 11 | 461 | 0.64 | 722 | 36.6 |
| 12 | 13 | 0.02 | 729 | 3.4 |
| 13 | 497 | 0.67 | 747 | 24.3 |
| 14 | 12 | 0.02 | 627 | 32.8 |
| 15 | 77 | 0.17 | 466 | 1.25 |
| Total<sup>†</sup> | 2871 |  | 7246 | Mean<sup>†</sup> 22.37 |
| *Patient 3 was excluded owing to high dropouts.<br>†Totals and means exclude Patient 3. |  |  |  |  |

We sought to investigate the relationship between critical slowing, epileptiform spikes (known biomarker of epilepsy) and seizures. We hypothesized that the likelihood of seizures would be related to the modulation of the autocorrelation and the variance signals. Furthermore, this modulation may also be linked to known changes in epileptiform activity [27, 28].

The variance and autocorrelation of the iEEG signals in each channel were computed in one second windows at two minute intervals for 14 of the 15 patients (patient 3 was not considered due to excessive data dropouts). The autocorrelation was considered to be the width at half maximum of the autocorrelation function (Figure 2B). The number of epileptiform spikes in each interval was also computed for each channel. Epileptiform spikes were detected using a previously described, patient-specific spike template and correlation algorithm [27]. Herein, we refer to *spikes* as the rate of epileptiform activity detected on each electrode.

It is known that seizures are often modulated by spikes and circadian and multidien (>24 hour) rhythms in a patient-specific manner [27, 28]. It is reasonable to suggest that critical slowing may also form a rhythmic process. To assess the temporal properties of the autocorrelation signal, a Fourier transform (FT) of the autocorrelation signal was computed. The peaks in the FT demonstrated that all patients had strong circadian rhythms (see Supplementary Figure 1). Some patients also had strong multidien rhythms with peaks between 3 and 30 days.

We investigated the relationship between seizures and the spike rates, autocorrelations, and variances derived from the iEEG on two temporal scales: short rhythms with periods of one day or less and long rhythms with periods with greater than two days. The long rhythms were extracted by applying a moving average filter with a length of 2 days to the signals. The short rhythms were extracted by subtracting the long rhythm from the raw signals and smoothing with a moving average filter of length 20 time-units. A Hilbert transform was applied to the signals to estimate magnitude and phase; the phase is important as it conveys whether a signal is increasing or decreasing at any given time. If critical slowing is occurring, we would expect that seizures occur with increasing probability when the autocorrelation and the variance signals are increasing (i.e. on the rising phase).

### Patient 1

Figure 2C shows the raw autocorrelation, variance and spike rate signals (gray), along with the long (orange) and short (dark blue) cycles of a sample channel for a representative patient, Patient 1. For clarity, the short cycle is shown prior to subtracting the long-cycle. Seizures are shown as red triangles. The coincident signal phases and seizure times were used to assess the relationships between seizures and the signals. Figure 2D illustrates the relationship between seizures and the long cycles. In each polar plot, the colored lines represent the normalized distribution of phases of the entire signal and the histogram represents the distribution of phases at the sample prior to the seizure times relative to the autocorrelation (gray), variance (red), and spike rate (cyan) signals. For this patient, seizures predominately occurred on the rising phases of the three signals and rarely occurred on the falling phases. Figure 2E similarly illustrates the relationship between seizures and the short cycles, also showing a predominance of seizures occurring on the rising phases of the signals, as expected from a process that is critically slowing.

Above each polar plot are the synchronization indices (SIs) for the signals (black) and for the seizure histograms (colored). The SI is a measure of phase uniformity in a signal (the tendency of a signal to have all phases uniformly distributed on a circle), or the synchrony between a rhythmic signal and events (seizures). A SI close to one indicates that a signal predominantly occurs at a single phase; a SI close to zero indicates that the signal is uniformly distributed across phases. For the seizure histograms, a SI close to one indicates that there is high synchrony between the phase of the signal and the times that seizures occurred. For Patient 1, the high SI across all measure demonstrated that there was a strong relationship between seizures and a given phase of the signal cycles.

### Critical slowing in 14 patients

Seizures tended to occur on the rising phase of the autocorrelation and variance signals as expected from a critically slowing process (Figure 3A; see also Supplementary Figures 2-15). In a few patients (Patients 8, 9, 11, and 13), many seizures unexpectedly occurred near the trough of the signals. The increased seizure rate near the trough of the autocorrelation and variance signals could be explained by two factors. Firstly, for these patients the autocorrelation and variance increased by a large amount through the transitions associated with the daily sleep-wake cycle, which is in itself a critical transition [24]. The change in autocorrelation was generally much larger than that caused by a seizure event itself. Seizures occurred on the trough of this much larger daily cycle. Secondly, most seizures were accompanied by large increases in spike rate prior to the seizure. Conversely, when the autocorrelation was high, there was a low spike rate (see Supplementary Figure 17 for two examples). It is likely that seizures in these patients were largely influenced by increases in spike rates [29], which could cause seizures during periods of low autocorrelation.

**Figure 3:**
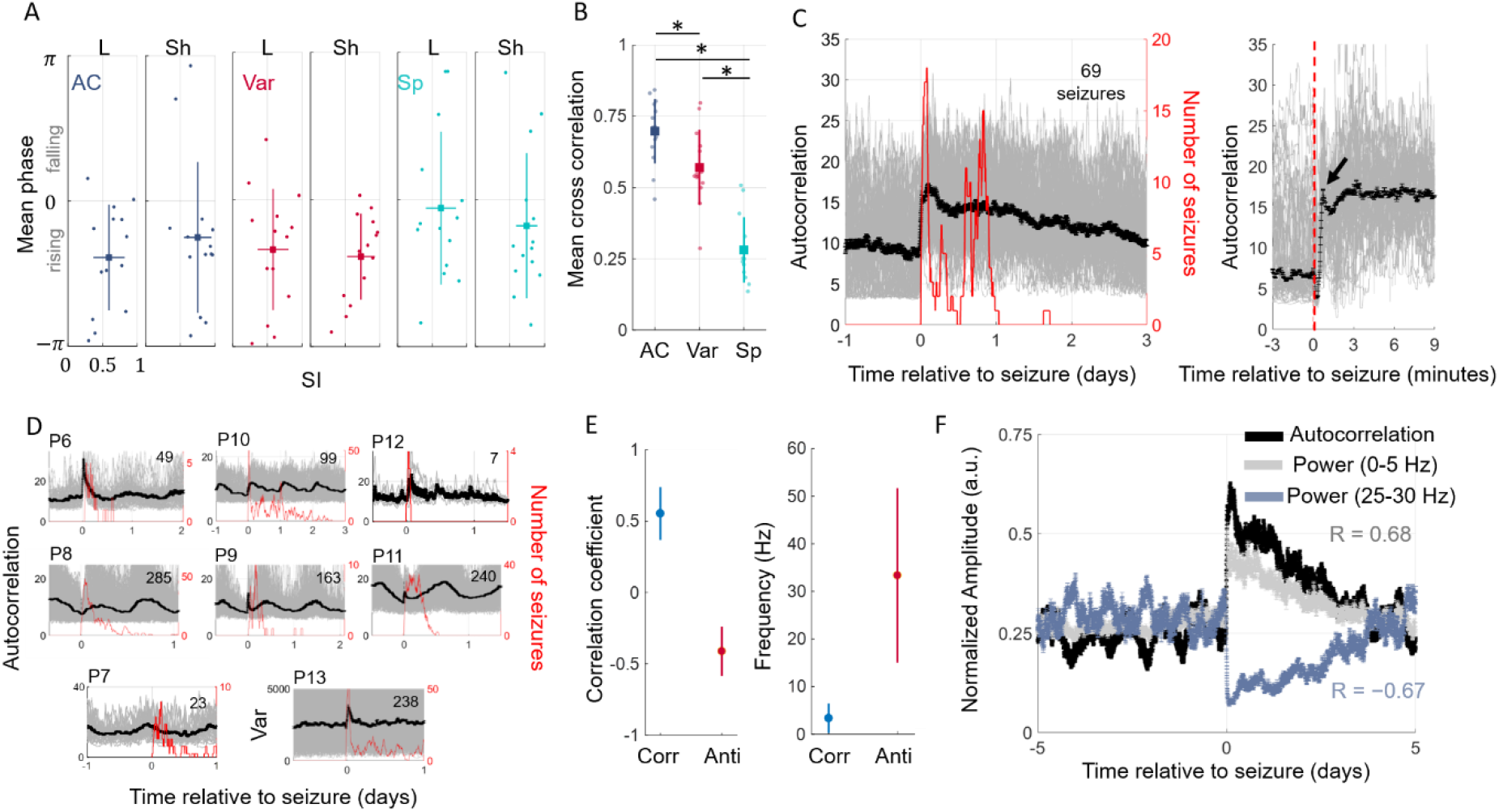
Critical slowing, seizure clusters and iEEG frequency. A) The synchronization indices (SIs) and mean phases at the sample prior to seizures for long (L) and short (Sh) cycles. A strong relationship between the three signals and seizures were observed as depicted by the high SI values. On average, seizures occurred on the rising phase of the signals. B) The similarity in autocorrelation (AC), variance (Var), and spike rate (Sp) signals across electrodes were compared by computing the mean cross correlation of each signal and comparing across patients. In A) and B), each dot represents a result for one patient; boxes indicate means across patients and bars indicate ±one standard deviation. C) Patient 1 had 69 seizure clusters. Individual seizures lasted a mean of 26 seconds, which is approximated by the arrow in the inset. Gray shows the raw autocorrelation from individual seizures. Inset represents an expanded plot of the autocorrelation computed on a finer temporal scale. D) Autocorrelation relative to the time from lead seizures for the remaining patients with seizure clusters. The number inset in each subplot denotes the number of lead seizures that occurred in clusters. For patient 13, the variance is shown instead of the autocorrelation. In C) and D), black lines denote the mean autocorrelation with standard error bars. Red line denotes a moving sum of seizures occurring after the lead seizure in a 2 hour window. E) Correlation coefficient between the iEEG power and the autocorrelation signal. Most patients showed a high positive correlation (Corr) between the autocorrelation signal and iEEG band power in the 0 – 10 Hz range. Many also showed a negative correlation (Anti) between signals in the 20 – 50 Hz range. Dots represent means and bars represent ±one standard deviation. F) Autocorrelation signal (black) and iEEG power in 0-5 Hz (gray) and 25-30 Hz (blue) frequency bands plotted relative to lead seizure times for Patient 1. Signals have been normalized for visual comparison. Lines represent means with standard error bars.

The SI was used to identify the electrodes that best captured the relationships between seizures and the underlying long and short cycles. The SIs across patients for long and short cycles are shown in Figure 3A. The SIs were greater than 0.5 for the short cycles for nearly all patients, suggesting a strong relationship between seizures and the short rhythms. Some patients also had a strong relationship between seizures and the long cycle.

Visual inspection of the autocorrelation signal in each patient showed that the signal tended to be similar across electrodes. Conversely, for the spike rate signal, the rates tended to be variable on each electrode. We quantified the similarity of the three signals (autocorrelation, variance, and spike rate) by computing a mean cross-correlation for the three signals independently across all channel pairs (Figure 3B). The average cross-correlation across all patients was 0.70, 0.57, and 0.27 for the autocorrelation, variance, and spike rate signals, respectively. There was a significant effect of signal type (F_2,13_ = 58.61, *p* = 2×10^-10^), and a post-hoc analysis showed that the autocorrelation signal was significantly different to the variance (*p* = 9×10^-3^) and the spike rate signals (*p* = 1× 10^-9^). This analysis suggests that the autocorrelation signal was consistent across the recorded brain areas and might be a measure that describes a state change throughout the brain, rather than a change in a localized brain region as is the case with spikes.

It is well recognized that seizures tend to cluster [30, 31]. We sought to examine the relationship between the autocorrelation and variance signals and seizure clusters. We found that nine of the fourteen patients had clusters of seizures that occurred within a short interval of a lead seizure. For a subset of these patients, the autocorrelation and variance signals slowly tended back to baseline after lead seizures over a period (hours to days) well beyond the duration of the individual seizures. During this time, there was an increased susceptibility to more seizures. Figure 3C shows an example of this relationship for Patient 1 (see also Figure 3C top row for Patients 6, 10, and 12). In this figure, lead seizures (gray) with a lead time >1 day are plotted along with a moving sum of seizures that occurred after lead seizures (red). The figure demonstrates how the autocorrelation slowly returned to baseline over a period of ~2 days and there was an increased susceptibility to seizures for ~1 day following the lead seizure.

For another subset of patients, both the autocorrelation and variance signals steadily increased after lead seizures. During this period there was an increased seizure susceptibility (Figure 3D middle row; see also Supplementary Figures for Patients 8, 9, 11), which was reduced when the autocorrelation and variance signals began to decrease.

From the perspective of critical slowing, these results are expected, as a high autocorrelation would suggest that the brain is in a prime condition for generating more seizures. The only exceptions were Patients 7 and 13. In Patient 7 there was an increase in seizure rate following lead seizures, despite there being a decrease in autocorrelation and almost no change in variance. For Patient 13, there was only an increase in variance and almost no change in autocorrelation (Figure 3D bottom row; see also Supplementary Figures for Patients 7 and 13).

iEEG signals are typically analyzed with respect to specific frequency bands that represent activity during different brain states [24]. It is possible that signals in a specific power band drive the cycles that we observed in the autocorrelation signal. To test this, we computed the iEEG band power and compared it to the autocorrelation signal. A high correlation was observed between the autocorrelation and iEEG signals in the 0-10 Hz band for most patients (Figure 3E, Corr).

Interestingly, a negative correlation was observed for signals in the 20 – 50 Hz band (Figure 3E, Anti). Although anti-correlated, it seems that also signals from this frequency band provides information highly relevant for the forecasting performance.

An example for Patient 1 is shown in Figure 3F. Here, the average autocorrelation signal relative to lead seizures (black) is shown against the average power in the 0 – 5 Hz range (gray) and 25 – 30 Hz range (blue). A high positive correlation was observed between the autocorrelation signal and the power in the 0-5 Hz range (R = 0.68) and a high negative correlation was observed between the autocorrelation and the power in the 25-30 Hz range (R = −0.67).

### Seizure forecasting

We evaluated the performance of a seizure forecasting algorithm based on the detected rhythms of the autocorrelation, variance, and spike rate signals. We consider two approaches:

- *Method M1*: Anti-causal filtering was used and potential forecasting performance was evaluated using all the available data. In this case, we evaluated the optimal level that a forecaster may perform using within-sample optimization, where prior knowledge of seizure susceptibility relative to the phase of the signals was taken from all the data. This method tests the performance of combining autocorrelation, variance, and spike rate signals in a forecaster and represents the best possible performance outcome of a forecaster.
- *Method M2:* Seizure rhythms were computed iteratively with a causal filter such that the forecaster for a given patient was based on information provided only by previous seizures. This approach tested out-of-sample forecasting performance in a pseudoprospective manner. Forecasting using this method began after the 10^th^ seizure. This method represents a forecaster based on the same signals but computed in a manner that is applicable to a clinical setting, where the algorithm learns as more data is available.

The relationships between seizures and signal phases were used to calculate the probability of a seizure. Figure 4A depicts the probability of a seizure occurring for Patient 1 using Method M1. Specifically, this is the probability of a seizure given the phases of the long and short cycles,

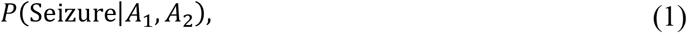

where *A*_1_ and *A*_2_ refer to the phases of the long and short cycles, respectively. From the probability distribution, the seizure probability versus time was calculated by multiplying the individual distributions under the assumption that each probability distribution was independent, since there was insufficient data to characterize the joint distributions accurately (Example for Patient 1 shown in Figure 4B, top). In an approach similar in nature to the original trial of the seizure advisory system that gave rise to the data considered here [21], our forecaster was designed such that, at any given time, a patient would be placed in a risk category: low, medium, or high risk. In practice, indicating the current risk level to a patient can be used to help guide their daily activities and encourage them to move to safety when seizure risk is high.

Two thresholds that optimally separated the low, medium, and high-risk categories were computed and used to categorize risk state over time (Example Figure 4B, bottom). The risk level assigned to each seizure is the risk state at the sample prior to the seizure occurring. For Patient 1 and using Method M1, 3% of seizures were assigned into low risk, and 90% into high risk categories. The proportion of time spent in the high-risk category was 3% and in the low risk category was 95%.

**Figure 4:**
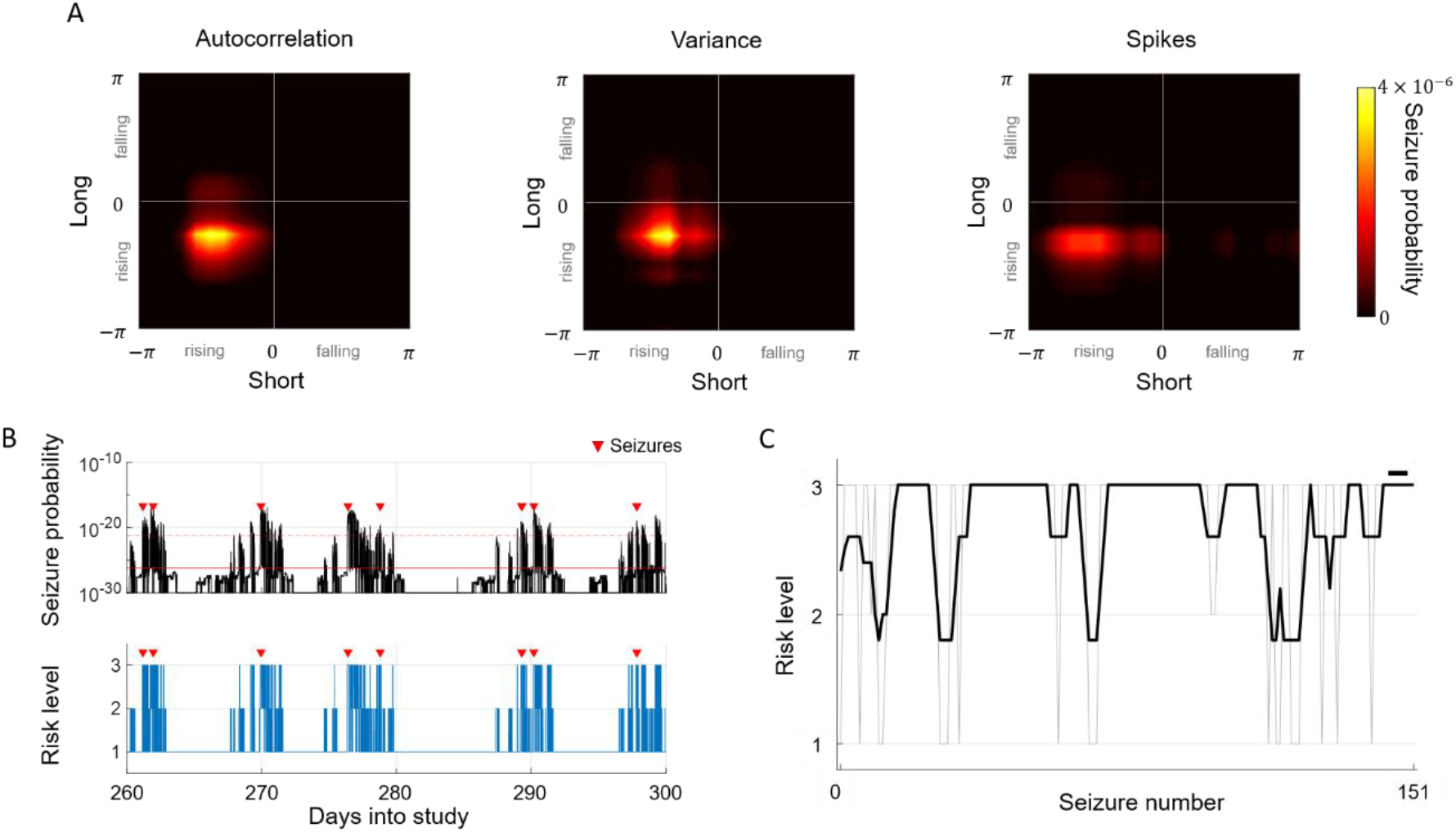
Seizure forecasting examples for Patient 1. A) The probability distributions of seizures given phases of the long and short cycles for the autocorrelation, variance, and spike rate signals. Brighter color corresponds to higher probability. B) The distributions from (A) were used to compute the probability of a seizure over time (top). Two thresholds separated low from medium risk (solid red) and medium from high risk (dashed red). The risk levels over time are shown in the bottom plot indicated by the heights of the blue bars; seizure times are indicated by red triangles. C) Pseudoprospective Method M2, where risk level at the time of each seizure is shown by the gray line and the black line denotes the five-seizure moving average risk (the black bar on the top-right denotes the length of the moving average).

To simulate a realistic situation that could be applied to clinical practice, we computed and updated the seizure probability distributions and risk levels iteratively after each new seizure (Method M2). Figure 4C shows the risk level assigned to each seizure (gray) and a moving average over five seizures (black) for Patient 1. The average risk level assigned to all seizures using Method M2 was 2.7. For Patient 1 and using Method M2, 15% of seizures occurred during low risk and 83% during high risk categories. The proportion of time spent in the high risk category was 8% and in the low risk category was 91%.

Figure 5A illustrates Method M2 approach applied to the data of the remaining patients (except for Patient 5 due to too few seizures). The average risk level assigned to seizures was 2.5 or greater for every patient, demonstrating that Method M2 achieved good performance. Figure 5B,C quantifies the performances of the two forecasters and compares it to a random predictor. There was a significant effect of method used to forecast seizures in the high risk category (F_2,12_ = 112.3, *p* = 7×10^-13^). A post-hoc analysis showed that both Methods M1 and M2 correctly classified significantly more seizures than the chance model (*p* = 1×10^-9^ and *p* = 1×10^-9^ for Methods M1 and M2, respectively). There were no significant differences between Methods M1 and M2 (*p* = 0.06). Furthermore, both Methods M1 and M2 had significantly fewer seizures in the low risk category than the random predictor (F_2,12_ = 148.3, *p* = 1×10^-9^ and *p* = 1×10^-9^ for Methods M1 and M2, respectively). There were no significant differences between Methods M1 and M2 (*p* = 0.5). No significant differences were found between the models in the amount of time spent in high and low risk states (F_2,12_ = 1.13, *p* = 0.3 and F_2,12_ = 2.12, *p* = 0.1 for high and low risk categories, respectively).

**Figure 5:**
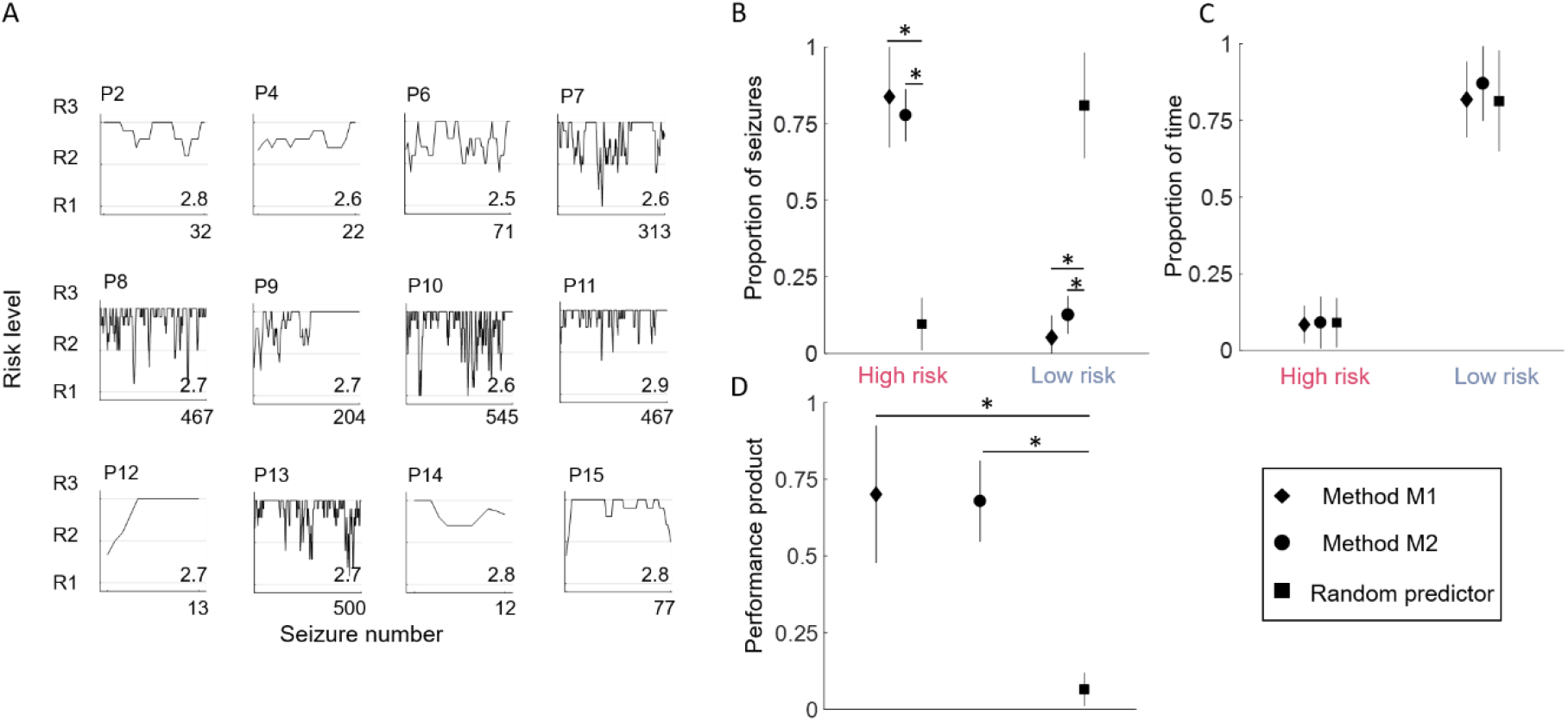
Forcasting performance for 14 patients. A) The pseudoprospective approach was applied to all patients, except Patient 5 due to too few seizures. A moving average (length 5 seizures) is shown for each patient. The number inset in each subplot represents the average risk level assigned to seizures. B) The proportions of seizures correctly classified as high risk or incorrectly classified as low risk are compared for Method M1 (diamond), Method M2 (circle), and a random predictor (square). C) The proportions of time spent in the high and low risk categories for the three methods. D) A performance measure, which was the product of the time spent in low risk, and the proportion of seizures classified as high risk, was used to compare the three models. In all plots, * represents significance to *p* < 0.01. The risk level for each seizure is taken as the risk level assigned to the sample prior to the seizure. Symbols in plots B, C, and D represent means and lines represent ±one standard deviation.

To compare the overall performances of the methods, the product of the proportion of seizures in the high-risk category and the proportion of time spent in the low risk category was used, which is hereafter called the “performance product”. In the ideal case, the performance product would be very close to one. Based on this measure, there was a significant effect of method (F_2,12_ = 73.7, *p* = 6× 10^-11^) and both Methods M1 and M2 performed significantly better than the chance predictor (*p* = 1×10^-9^ and *p* = 5×10^-9^ for Methods M1 and M2, respectively; Figure 5D). There was no significant difference between Methods M1 and M2 (*p* = 0.2).

Seizure forecasting performance was compared using measures from critical slowing versus spike rates alone, and then the combined measures (Supplementary Figure 16). The performance product was used to compare the three different cases. There was a significant effect of method on performance (F_2,12_ = 8.9; *p* = 0.0013), and the combed approach performed significantly better than using spikes alone (*p* = 9× 10^-4^). Seizure forecasting performance was also compared to the performance of the original trial ([21]; Supplementary Figure 18). Method M1 achieved a higher sensitivity and specificity overall compared to Method M2 and the original trial. Finally, we compared our pseudoprospective forecasting results to other pseudoprospective algorithms previously developed on the same dataset. This included a machine learning algorithm [32], a predictor based on circadian rhythms and logistic regression [33], and multiple algorithms based on a crowd-sourced approach to seizure prediction [34]. Method M2 scored higher on sensitivity (seizures in high risk), and a lower time-in-high for all patients except Patients 7 and 14 (Supplementary Figure 19).

## 3. Discussion

Critical slowing is a marker of decreased resilience as a system approaches a tipping point. Prior to this study, critical slowing had been theorized to occur in the brain as seizure susceptibility increased [3, 14, 16, 18]. However, there has been little evidence of critical slowing prior to seizures demonstrated in humans. We previously demonstrated that features of critical slowing are present prior to seizures in work focused on *in vitro* and *in vivo* models [35]. Furthermore, interictal spikes were shown to have both a seizure generating, or seizure suppressive effect depending on when they occur relative to the dynamical state of the brain. The current paper is a unifying progression where we have investigated the interplay between critical slowing, long-term rhythms, interictal spikes and seizures in humans.

In the present study we have shown that two features indicating the presence of critical slowing – autocorrelation and variance – increase prior to seizures and are representative of seizure susceptibility. These signals were combined with epileptiform spikes to create a powerful forecasting tool. The long-term and continuous nature of the data used in this study was extremely important, as signal changes occurred over much longer time scales than had previously been assumed in models of seizure onset. The long, patient-specific time-scales also explain the ambiguity in the previous literature, which typically used shorter, fixed windows to detect critical slowing [18].

Our work contributes four major findings to the field:

1. Evidence that seizures are preceded by critical slowing as suggested by computational models of epilepsy. This provides important validation of the mathematical models used in epilepsy that have so far proved difficult to apply in humans.
2. Markers of critical slowing were not confined to a localised region in the brain, suggesting that changes in susceptibility are detectable across broad areas of the brain.
3. Seizures tended to occur on a narrow phase of the periodic autocorrelation, variance and spike rate signals. These signals provide a powerful forecasting tool that can be used to determine seizure susceptibility.
4. Seizures tended to cluster after lead seizures. The period during clusters was also characterized by a sustained increase in the autocorrelation, which is consistent with critical slowing. This result provides a mechanistic basis for seizure clusters.

Critical slowing has been observed prior to seizures in experimental studies [16, 35, 36] and at the end of most seizures in humans [13]. Figure 1A illustrates how a bifurcation can lead to critical slowing in the EEG signal. While other bifurcations are also plausible, noise induced fluctuations are expected to increase in intensity near any critical (i.e., second or higher order) phase transition and display the characteristic features of critical slowing [16]. The example in Figure 1A may be oversimplified, but it captures an essential aspect of the dynamical changes that are supported by our results. The example demonstrates how seizure susceptibility can change on a slow temporal scale by variations in parameter *k*, and how noise induced fluctuations can trigger a seizure in a probabilistic manner [3, 35].

Milanowski et al. [18] found that, in most patients, brain activity recorded via iEEG did not show signs of critical slowing leading up to a seizure. Our results do not contradict their observations, but instead demonstrate that the warning signals fluctuate over longer temporal scales (hours and days) than those considered in their study (seconds to minutes). Indeed, there is accumulating evidence that suggests seizures are mediated by long cycles [37]. Seizures and subclinical epileptic activity, such as spike discharges, have been found to be distributed into circadian cycles [27, 30]. More recently Baud et al. [28] and Karoly et al. [38] found that epileptiform activity and seizures fluctuate with daily and multi-day rhythms. The current study builds on these previous analyses by showing that seizures arise as the brain approaches a critical transition, and that the autocorrelation and variance signals fluctuate with similar rhythms to those found in epileptiform activity.

For most patients, both the autocorrelation and variance signals remained high for a prolonged time after a lead seizure, far longer than the duration of the seizure itself. This led to a state where seizures were more likely to keep occurring, leading to seizure clusters. The result was unexpected, as seizure termination has been suggested to follow a critical transition, leading to a low variance post-ictal state [13]. It should be noted, however, that this previous work explored a much shorter post-ictal window than considered in the current paper. While the mechanisms for seizure clusters are poorly understood, it has been suggested that an ictal focus becomes more excitable, or less inhibited following a first seizure [30, 31]. This observation is supported by our results, where a high autocorrelation and variance following a lead seizure would suggest that the brain remains in a highly susceptible state.

In this study, two methods for seizure forecasting were explored: Method M1, where seizure rhythms were calculated using all available data, and Method M2, where seizure probabilities were iteratively computed based on past seizure occurrences. Both methods could accurately forecast seizures (sensitivity 84±16% and 77±8%, respectively, when averaged across patients), performing significantly better than chance (9.6%). In the original study [21], 72±13% of seizures were correctly classified as high risk during the training phase. However, the performance dropped to 58±25% during the advisory phase (Supplementary Figure 18). The percent of time spent in high risk in the original study was greater than in the current study: 31±8% and 25±10% during the training and advisory phases, respectively, c.f. 8±6% and 9±8% using Methods M1 and M2, respectively. It should be noted that the original trial used only a subsample of the data used in the current study.

Seizures occurred in a patient-specific and probabilistic manner relative to the two warning signals (i.e. seizures did not always occur when the signal variance and autocorrelation increased). Forecast performance significantly improved when the combination of autocorrelation, variance and spike information was included compared to using spike rates alone (see Supplementary Figure 16), demonstrating the power of using additional predictors. The need for combining statistical priors is a framework that is being accepted in the seizure forecasting community and new methods for combining multiple predictors are emerging [2, 34]. The future of seizure forecasting will undoubtedly include multi-modal information, combining a patient-specific mixture of implanted and wearable technologies [1, 2, 39].

Multimodal information will be particularly important for differentiating between different types of state transitions. In a bistable, or multistable system, an increasing autocorrelation is indicative of an approach to a critical transition, from which a state change can occur. The state change in the human brain could be from the wake/sleep cycles, anaesthesia [40], depression [9] or seizures. Indeed, we found that the sleep cycle produced large changes in the autocorrelation and variance which could have confounded the results, especially for patients who had a larger seizure probability during the trough of the daily cycle (see Supplementary Figure 17). Differentiating between the types of possible brain states, their interactions and associated complexity is an important future endeavor.

The underlying mechanisms that modulate seizure susceptibility remain unknown. A few patients had ~12 or ~24-hour cycles, suggesting an influence of circadian rhythms. These rhythms could be influenced by hormonal fluctuations, such as changes in cortisol and melatonin [41, 42].

Anti-epileptic drugs play an important role in modulating cortical excitability [43] and, thus, likely influence the patterns observed in this study. However, Patient 6 was not on medication, yet had a strong circadian cycle, demonstrating that the circadian influence on seizures was not modulated by anti-epileptic drugs in this patient. The multidien cycles observed for most patients highlight the presence of slow variables (>24 hours) that influence seizure susceptibility [28, 38, 44]. The causal factors of these slower cycles may be regulated by the body [45] or relate to external factors such as weather [46] or behavior [47]. Identification of the causal factors will undoubtedly improve our understanding of seizures and improve techniques that make seizures predictable.

It is clear from this, and other recent studies [28, 38] that long-term monitoring in epilepsy will be necessary to create patient specific clinical treatments. The autocorrelation signal was surprisingly very consistent across electrodes (Figure 3B), suggesting that this signal may be an effective measure to monitor state changes in the brain. The autocorrelation signal was best represented by the lower frequency power band of the iEEG (0 – 10 Hz). For many patients, it was anti-correlated with signals in the 20 – 50 Hz band, which may also provide important forecasting information.

There were two limitations in the work presented here. Firstly, the signals used in this study were wideband signals. The Hilbert transform is typically computed on narrow-band signals and can be ill-defined for wideband signals. This is because wideband signals can be represented as the superposition of smaller signals with multiple frequencies. To mitigate this effect, we separated the time series signals into two separate temporal domains by smoothing with a 2-day window (long cycle) and one with a 20 sample (40 minute) window (short cycle). It is possible that the phase estimation would be improved by smoothing further. However, this may come at a cost of reduced forecasting performance, as quick changes in the signals being tracked would be less obvious.

Another limitation was that many seizures did not occur at the peak of the phase cycle (highest seizure probability) as expected from critical slowing; rather they occurred just prior to the peak (i.e. Figure 2D,E). This is most likely because the seizure event itself is characterized by an even larger increase in autocorrelation and variance, which confounds the analysis. Hence the peak in the autocorrelation occurs after the seizure event begins. This is most obvious in Patients 4 and 6 (black arrow in Supplementary Figures 4 and 6), where seizures were preceded by a small increase in autocorrelation, but the seizure event itself resulted in a very large increase that lasted several hours.

In a couple of cases (Patient 8, 9 and 11), seizures often occurred near the trough of the autocorrelation cycle, which is contrary to critical slowing as we expect there to be the lowest seizure probability. It is possible that critical slowing does not precede seizures in these patients. However, the metrics used in critical slowing were still very useful for seizure forecasting. It is interesting to note that in these patients, increases in autocorrelation were accompanied by low spike rates, and periods of low autocorrelation were characterized by very large increases in spike rate (Supplementary Figure 17). It is possible that in these patients, interictal spikes were the main contributors to seizure genesis.

Seizure forecasting has the potential to transform the clinical approach to the treatment of epilepsy. An accurate forecast could be used to provide the patient a warning and also trigger interventions. Here, we have applied a theoretical approach, critical slowing, to the forecasting of seizures. This approach outperformed all previous attempts to predict seizures on the same dataset (Supplementary Figure 19). It is important to note that the time in high risk was defined differently for the various studies, hence they may not be directly comparable. Our approach has been designed to give a warning at the sample prior to a seizure (i.e. at least 2 minutes prior to a seizure). A clinical device could be used to intervene during periods of high risk by, for example, applying deep brain stimulation as required. Future interventions that incorporate forecasting will undoubtedly pave the way towards improved outcomes for people living with epilepsy.

## Supporting information

Supplementary figures summarizing the results for each patient, and the forecasting performance against other previously developed algorithms

## Acknowledgements

IEEGPORTAL and National Health and Medical Research Council Project Grant 1130468.

CM acknowledges support from a NARSAD Young Investigator grant.

MM acknowledges support from the Melbourne Neuroscience Institute Fellowship award.

DRF, LK, and MC acknowledge the support from the Epilepsy Foundation of America, My Seizure Gauge Challenge Funding. PJ acknowledges support from the Ministry of Health of the Czech Republic 17-28427A.

## 4. Online Methods

A total of 15 patients with focal epilepsy participated in the first-in-human study (for patient details, see Cook et al. [21]). All data were collected with ethics approval from Human Research Ethics Committees at the participating institutes. The seizure advisory system captured continuous iEEG recordings on 16 electrodes at 400 Hz sampling rate (Figure 2A). Patients were implanted with 16 electrodes placed near the presumed epileptogenic zone. A board-approved epileptologist reviewed each patient’s iEEG and annotated the seizures and their durations. Public access to a subset of this data is now available on the online platform Epilepsyecosystem [34]. Spikes in the data were detected using a correlation-based algorithm that compared the iEEG signal to a patient-specific template [27]. Example spikes for Patient 1 are shown in Figure 6A.

**Figure 6:**
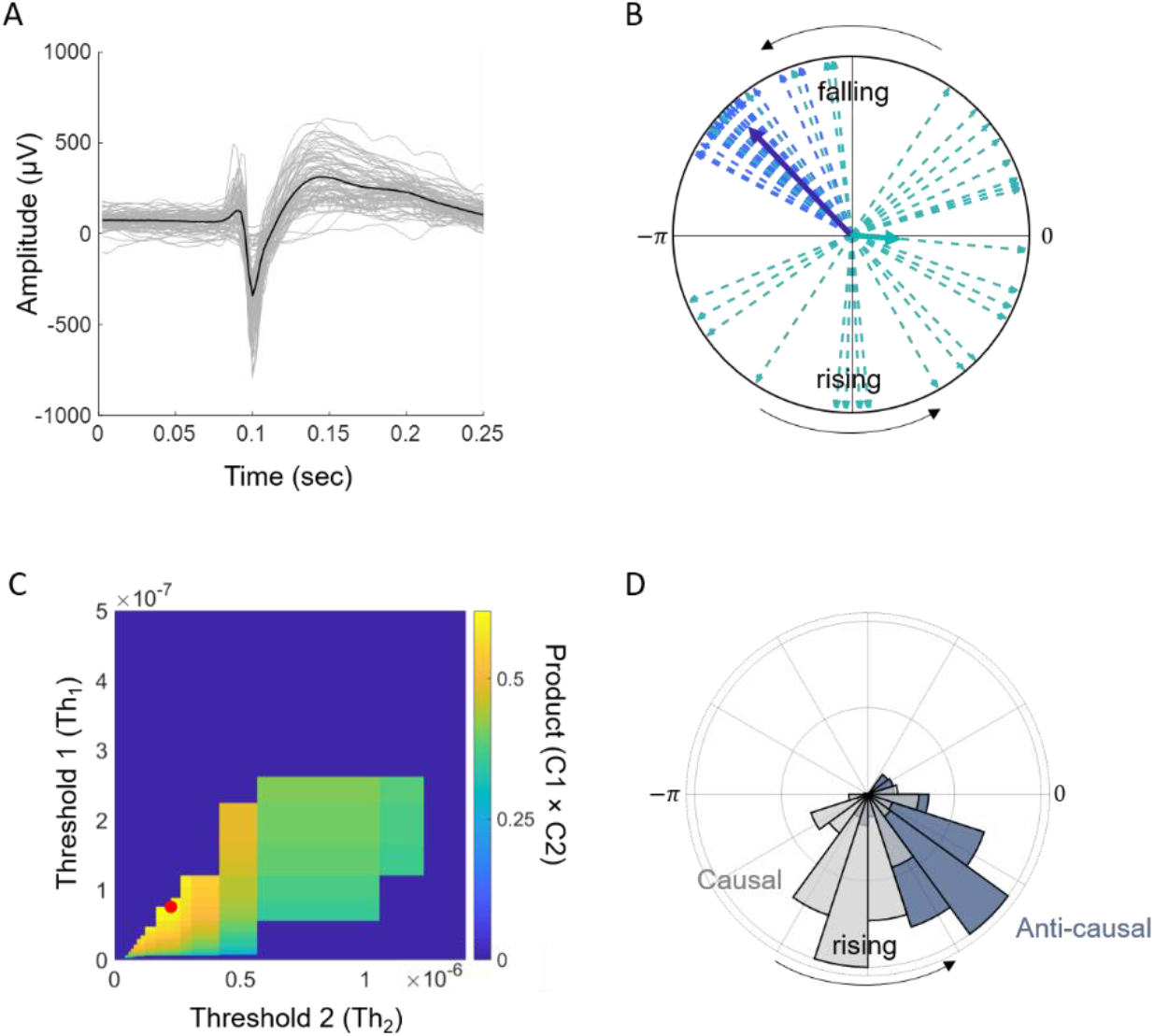
Methods used in the analyses. A) Example of epileptiform discharges (termed ‘spikes’ throughout the text) detected using a patient-specific template matching algorithm. B) The synchronization index (SI) is computed by adding complex phases (dashed vectors) and computing the mean resultant length (bold vectors). Shown are two examples, one with a high SI (dark blue) and one with a low SI (green). C) Two thresholds were optimized to separate seizure probability into low, medium, and high risk states by maximizing the product of criteria C1 and C2. Regions where criteria C3 and C4 were not achieved were set to zero. In this example, the red dot shows the optimal thresholds. D) An example of the phase shift observed when using causal (light gray) or anti-causal (dark gray) filtering.

### Data selection and pre-processing

All analysis and data manipulation were conducted using MATLAB (MathWorks 2017a). iEEG recordings were divided into one-second segments separated by two minutes (Figure 2A). In each segment, the signal variance (Equation (2)) and autocorrelation (Equation (3)) were computed,

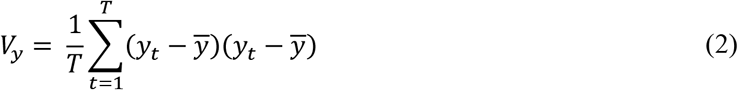

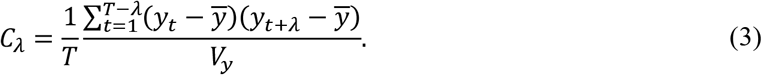

Here, *T* represents the number of samples in signal *y* and *ȳ* represents the signal mean. *C_λ_* represents the autocorrelation function of signal *y* as a function of the lag value *λ*. The autocorrelation measure used in the study was taken as the width at the half maximum of the autocorrelation function (Figure 2B). We also investigated the lag 1 autocorrelation measure [18] and found that it gave approximately similar results. However, the autocorrelation width produced a larger dynamic range of values from which to observe changes in the signal.

After computing the autocorrelation, variance, and spike rates for the entire dataset, a moving average filter was applied with a window of 2 days (1440 samples) to identify long rhythms in the data. The filtered data was then subtracted from the unfiltered data to identify short rhythms. This signal was then smoothed using a moving average filter of length 20 samples (40 min). A Hilbert transform (MATLAB’s *hilbert* function) was applied to the long and short rhythms to compute the analytic signal, from which signal phases could be derived.

The autocorrelation and variance signals were also computed in one second windows with no separation between windows. These were computed 3 hours prior to and after each seizure (i.e. Figure 3C, inset). The data was used to investigate seizure susceptibility after a lead seizure in finer detail.

### Intracranial EEG (iEEG) frequency

The autocorrelation signal was compared to the iEEG power in specific frequency bands. The Chronux toolbox for MATLAB (http://chronux.org/), with 5 tapers and a time bandwidth of 3 was used to estimate the spectral power for each one second segment of the iEEG. The spectrum was then summed in bins of 5 Hz to obtain the power in each 5 Hz segment. This produced 20 time-series for each 5 Hz segment from 0 to 100 Hz. The time series had equal length to the autocorrelation, variance and spike rate signals. These power signals were smoothed using a moving average filter of length 20 (similar to the autocorrelation, variance and spike rate signals) and then compared to the autocorrelation signal. Correlation coefficients were computed by first determining the correlation between the power in each 5 Hz segment and the autocorrelation signal. This resulted in a maximum positive and negative correlation for a particular 5 Hz band in each patient, which is reported in Figure 3E.

### Missing data

Over the course of the study, many data dropouts occurred when the recording system was not fully recharged or when data was not regularly retrieved. As a result, all patients had gaps in their data, lasting from minutes to days. In most cases, the dropouts were short segments. In one case (Patient 3), the segments of dropout data accounted for almost 80% of the total recording duration, so this patient was removed from analysis. For the remaining patients, the autocorrelation, variance, and spike rate were computed for each electrode.

Short sections (<2 hours) in the autocorrelation, variance, spike rate and power signals that contained dropouts were filled with Gaussian noise. The mean and standard deviation of the noise was computed from the remaining data without dropouts. Sections with larger gaps were left as missing values. These missing values were ignored when computing averages.

When computing the Hilbert and the Fourier Transform (FT), missing data were first filled with Gaussian noise (as with the shorter sections). This has the effect of introducing noise into the FT and the Hilbert transform. When computing the signal phases from the analytic signal, 60 samples (2 hours) either side of dropouts were removed from analyses to reduce the effect of boundaries.

### Synchronization index

The phases at the times of the seizures (Figure 2C, red triangles) were used to calculate the synchronization index (SI) [48]. Each phase, given by the analytic signal derived from the Hilbert transform, is represented by a complex number that can be drawn on polar axes as a vector (Figure 6B, thin arrows). The SI is given by

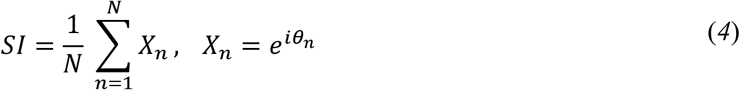

where *X_n_* represent the complex valued analytic signal of the autocorrelation, variance, or spike rate signals at the sample number *n*, and *θ_n_* is the phase of the signal. 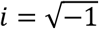 and *N* represents either the total number of seizures, or the length of the signal depending whether the SI was being used to compute the seizure histograms, or the phase uniformity of the signal (Figure 2D,E). If all seizures occur at nearly the same phase of the filtered signal, then the SI will be close to 1 (e.g. dark blue vector in Figure 6B). If seizures occurred on random phases of the filtered signal, the SI will be close to 0 (e.g., the green vector in Figure 6B). The SIs reported in this study were computed the sample prior to each seizure.

### Similarity between electrodes

The similarities between the three signals across electrodes were compared using a correlation coefficient. The autocorrelation, variance, and spike rate signals were first smoothed using a moving average filter of length 20 and sections containing missing values were removed. A cross-correlation (MATLAB corrcoef function) was then computed between the signals on each electrode. The cross-correlation produces a 16×16 matrix representing the correlation between each electrode combination. The matrix was transformed into a vector with the diagonal and duplicate values excluded, and the mean correlation was then computed and compared (Figure 3A).

### Seizure clusters

Seizure clusters were determined by analyzing the inter-seizure intervals for each patient. We plotted a histogram of seizure intervals with a bin spacing of 1 hour. Patients that did not have a seizure count of at least 5 within the first day were not considered to have seizure clusters. For the remaining patients, we used the histogram to determine the seizure lead times. The histograms showed two types of responses: (1) there was an exponential decay from time zero or (2), there were multiple peaks at regular intervals. For example, Patient 1 had a peak close to zero and an exponential decay without any other obvious peaks. Patient 9 had multiple peaks at daily intervals (see Supplementary Figures for Patients 1 and 9). For cases where there was an exponential decay, we set the lead time to be 1 day. For patients where there were multiple peaks, we set the lead time to the trough between peaks. For Patient 9, a lead time of 0.6 days, or 14.4 hours was chosen. Seizure clusters were investigated relative to the autocorrelation and variance signals on the channel with the highest SI.

### Seizure forecasting

Seizure risk was computed for each patient using a probability distribution of seizures relative to signal phase. The resulting probability density was used to compute seizure probability over time, from which three risk levels were determined: low risk, medium risk, and high risk. The risk levels for were computed using thresholds on the seizure probability that were optimized to achieve the following criteria:

C1. Maximize the time spent in low risk periods.
C2. Maximize the number of seizures classified in high risk periods.
C3. The time spent in low risk is greater than time spent in medium risk. The time spent in medium risk is greater than time spent in high risk
C4. The number of seizures occurring during low risk is less than the number occurring during medium risk. The number of seizures occurring during medium risk is lower than the number occurring during high risk.

The optimization was conducted by maximizing the product of C1 and C2. For Method M1, the product of C1 and C2 at points where C3 and C4 were not satisfied was set to zero. For Method M2, setting these points to zero often resulted in no optimal solution being found, hence optimization was only conducted on C1 and C2. Since the search space was small, the thresholds could be optimized quickly using a brute-force approach. Figure 6C shows the search space for Patient 1. Threshold 1

(Th_1_) corresponds to the threshold separating the low and medium risk states. Threshold 2 (Th2) separates the medium and high risk states. The combinations of thresholds that best achieved the above criteria are shown by a red circle.

### Method M1

The potential to forecast seizures was evaluated using the short and long rhythms of all three signals (autocorrelation, variance, and spike rate) over all the data. The probability of a seizure given phase was computed using the phase estimated from the analytic signal. The phases between — *π* ≤ *θ* < *π* were broken up into 20 equally spaced windows. A probability given phase was computed by evaluating the number of seizures that occurred in a phase window (*S_θ_*) divided by the number of times the phase appeared in the signal (*N_θ_*):

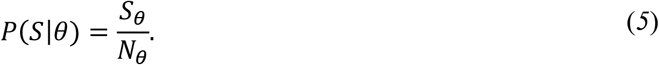

The probability density for the combined short and long rhythms, and for the combined autocorrelation, variance and spike rate signals were computed by multiplying the probabilities together.

### Pseudo-prospective seizure forecast (Method M2)

The advantage of using a moving average filter to identify the short and long rhythms is that, when it is applied causally, it introduces a predictable phase shift in the data (Figure 6D). The relationships between seizures and the signals remain largely unchanged apart from a constant phase shift. Forecasts using Method M2 employed causal filtering to identify the short and long rhythms, and to estimate the phase relationships between seizures and the signals. The risk level for each seizure was determined iteratively using the seizure rhythms and risk level thresholds that were determined from past seizure information bootstrapped using data from the first 10 seizures. When a new seizure occurred, the relationship between seizures and the signal phases were recalculated and used to re-estimate the seizure probability distribution, which remained fixed until the next seizure. Due to non-stationary effects in the signals [49], the seizure probability distribution was calculated over a 50 day window.

### Random predictor

The performance of the two forecasting methods were compared to a random predictor using a random Markov model. To compute the random model, the transition probabilities for each risk state were first computed (based on Method M1); i.e., the probability of transition from low risk to medium risk, medium to high risk etc. Then, a model that randomly transitioned between the three risk states was then generated using the transition probabilities. Statistical differences in the data were computed using ANOVA followed by post-hoc analysis using Tukey-Kramer comparison, where necessary, with alpha = 0.01.

