## Supplementary figures summarizing the results for each patient, and the forecasting performance against other previously developed algorithms for "Critical slowing as a biomarker for seizure susceptibility"

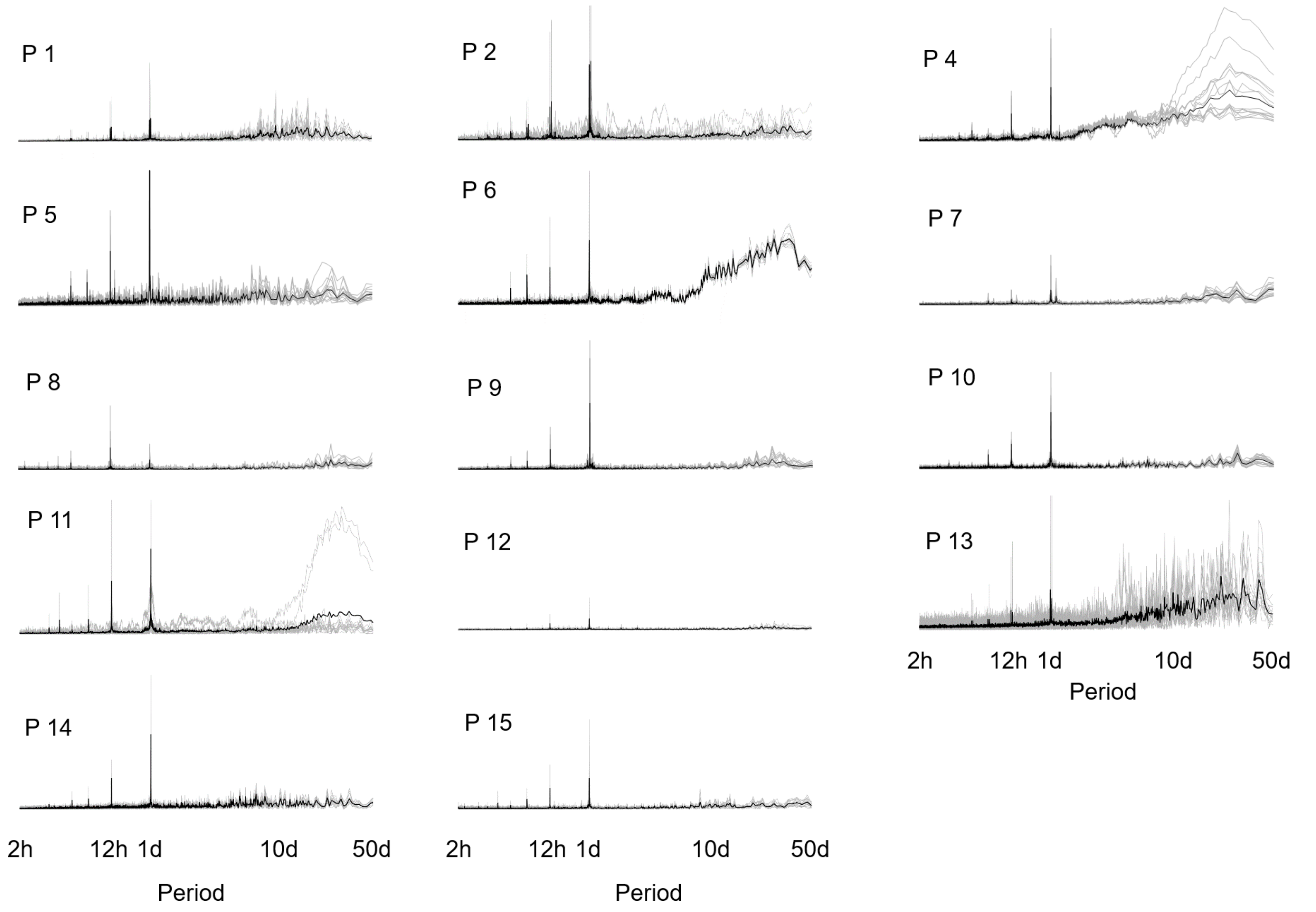


Supplementary Figure 1: Fourier transform (FT) of the raw autocorrelation signal for each patient. The transform shows peaks at 12 and 24 hours for every patient, demonstrating a strong daily rhythm in the signal. Some patients also had peaks at higher periods (e.g. Patients 4 and 6). In each subplot, the gray lines represent the FT for each of the sixteen channels, and the black represents the mean across channels.

### Patient 1

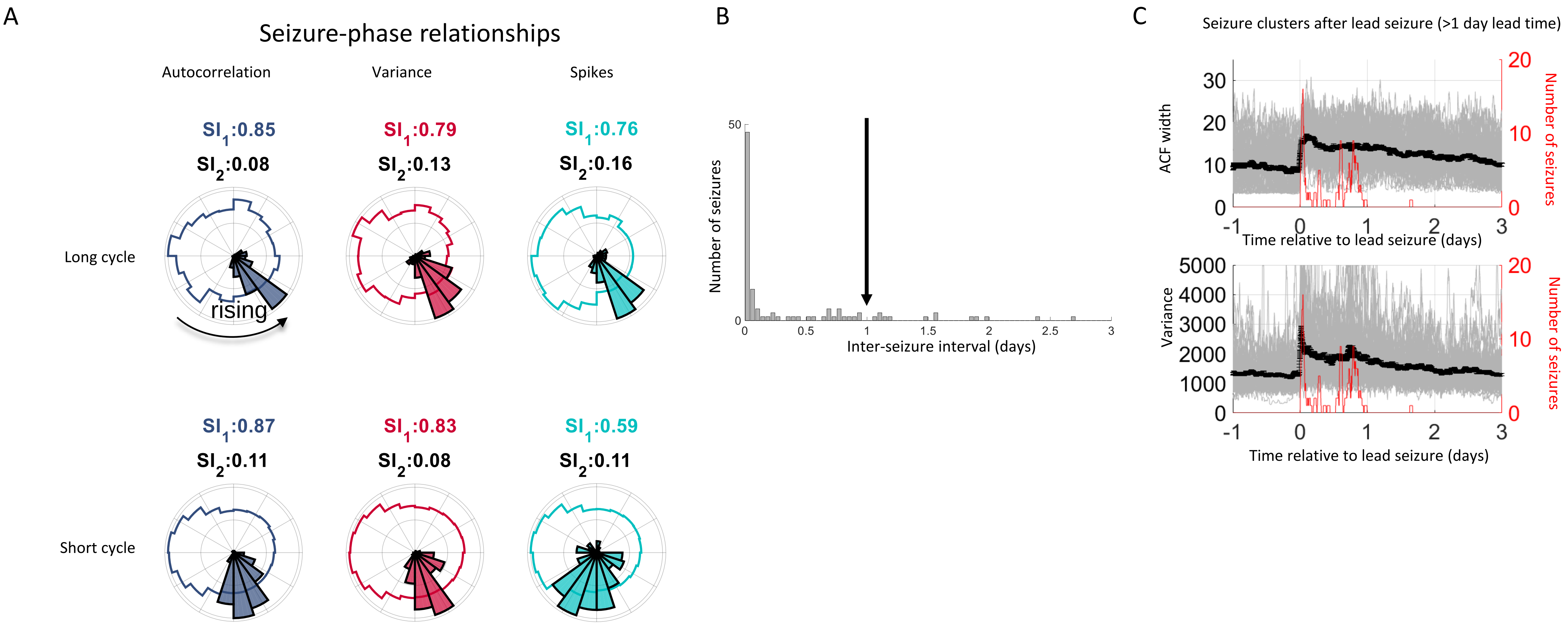


Supplementary Figure 2: Results summary for Patient 1. A) The Synchronization Indices ($SI_1$) for the autocorrelation (gray-blue), variance (red) and spikes (cyan) quantify the synchrony between seizures and the underlying signals. A high $SI_1$ corresponds to good alignment between seizures and the phase of the signal. The SI for the signals ($SI_2$) quantifies the phase uniformity of the signal. A low $SI_2$ demonstrates that the signal is periodic and that all phases are equally represented. B) The inter-seizure interval showed that a very high proportion of seizures occurred within 1 hour of a previous seizure. This patient had an exponential decay in the histogram. We set the lead seizure cut-off at 1 day. C) Lead seizures (gray) are plotted relative to the time of the seizure. After a lead seizure, the autocorrelation and variance signals decayed to baseline over a few days. During this period, there was an increase susceptibility to seizures. The black lines shows the average with standard error bars. Red line shows a moving sum of seizures in a 2 hour window.

### Patient 2

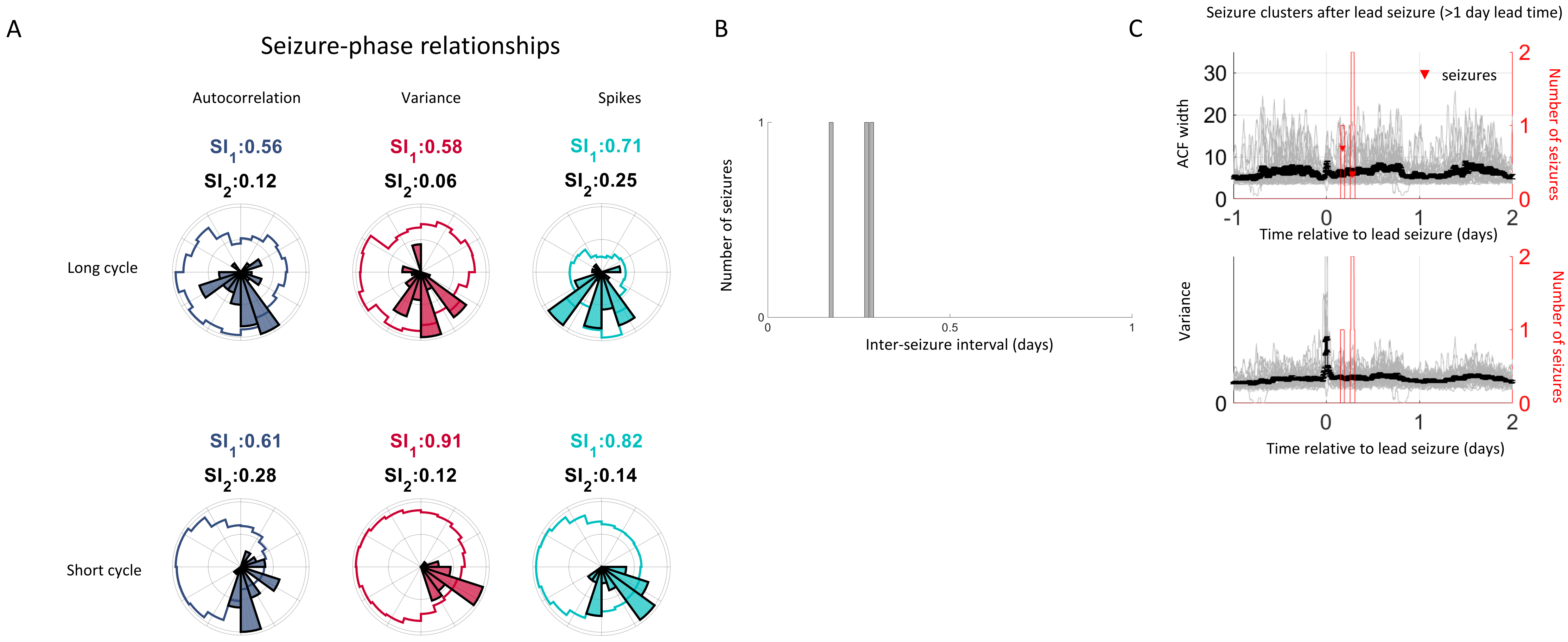


Supplementary Figure 3: Results summary for Patient 2. A) The Synchronization Indices ($SI_1$) for the autocorrelation (gray-blue), variance (red) and spikes (cyan) quantify the synchrony between seizures and the underlying signals. A high $SI_1$ corresponds to good alignment between seizures and the phase of the signal. The SI for the signals ($SI_2$) quantifies the phase uniformity of the signal. A low $SI_2$ demonstrates that the signal is periodic and that all phases are equally represented. B) The inter-seizure interval. This patient had too few seizures to meet the seizure clustering criteria. C) Lead seizures with a lead time of 1 day (gray) are plotted relative to the time of the seizure. The autocorrelation and variance did not tend to stay elevated after a lead seizure. The black lines shows the average with standard error bars. Red line shows a moving sum of seizures in a 2 hour window.

### Patient 4

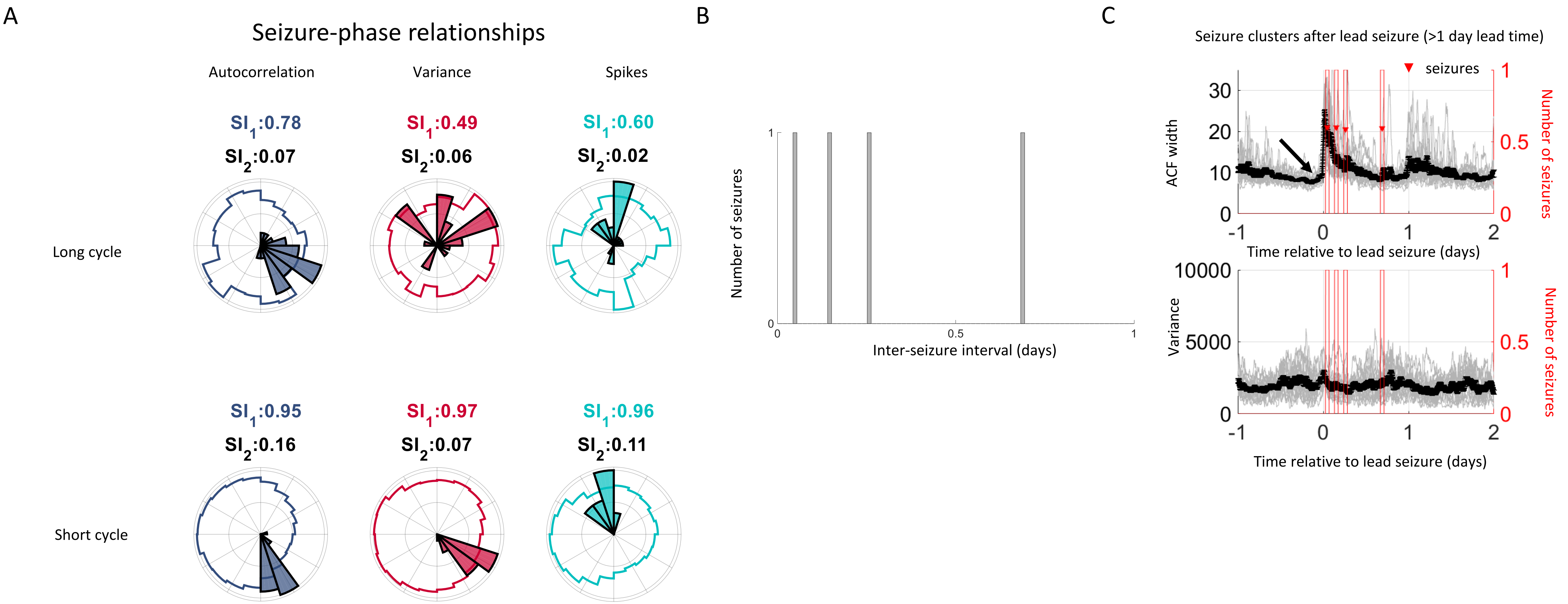


Supplementary Figure 4: Results summary for Patient 4. A) The Synchronization Indices ($SI_1$) for the autocorrelation (gray-blue), variance (red) and spikes (cyan) quantify the synchrony between seizures and the underlying signals. A high $SI_1$ corresponds to good alignment between seizures and the phase of the signal. The SI for the signals ($SI_2$) quantifies the phase uniformity of the signal. A low $SI_2$ demonstrates that the signal is periodic and that all phases are equally represented. B) The inter-seizure interval. This patient had too few seizures to meet the seizure clustering criteria. C) Lead seizures with a lead time of 1 day (gray) are plotted relative to the time of the seizure. The arrow denotes a visible increase in autocorrelation prior to the seizure that is overshadowed by the seizure event itself. The autocorrelation decayed back to baseline over approximately 12 hours, during which time the patient had three seizures. The variance signal did not show the same pattern. The black lines shows the average with standard error bars. Red line shows a moving sum of seizures in a 2 hour window.

### Patient 5

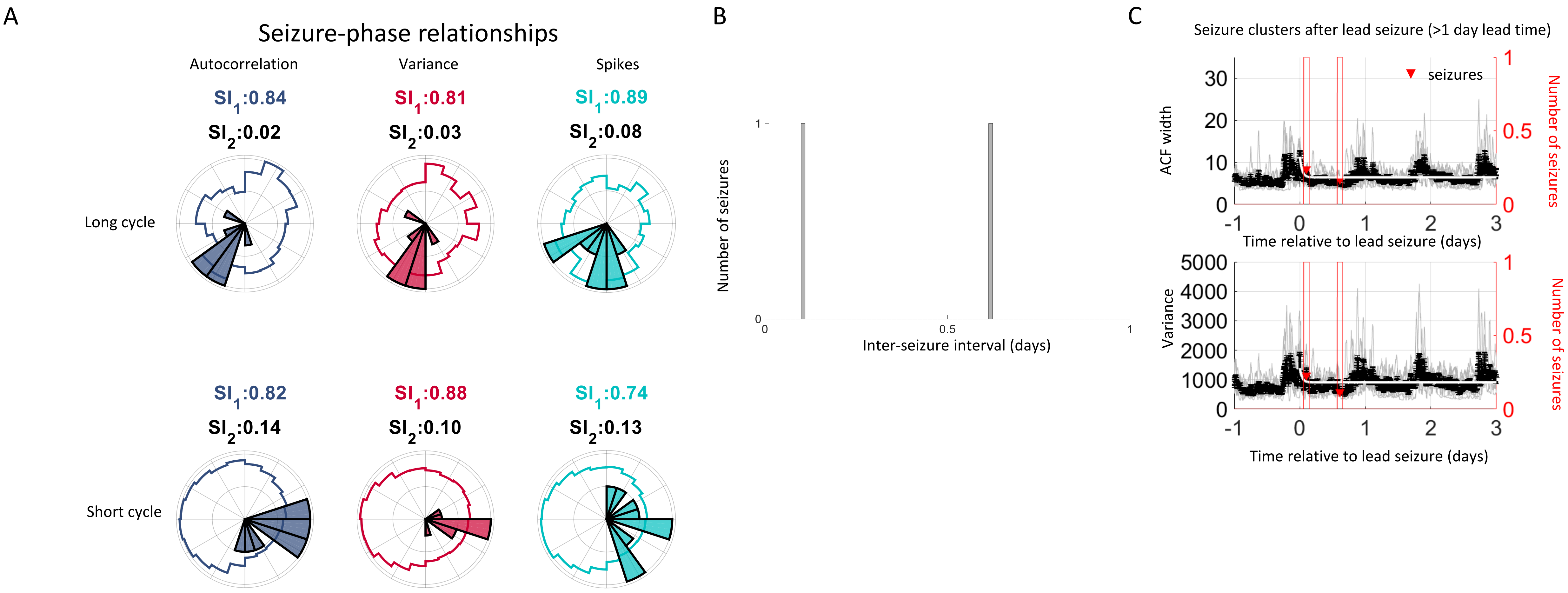


Supplementary Figure 5: Results summary for Patient 5. A) The Synchronization Indices ($SI_1$) for the autocorrelation (gray-blue), variance (red) and spikes (cyan) quantify the synchrony between seizures and the underlying signals. A high $SI_1$ corresponds to good alignment between seizures and the phase of the signal. The SI for the signals ($SI_2$) quantifies the phase uniformity of the signal. A low $SI_2$ demonstrates that the signal is periodic and that all phases are equally represented. B) The inter-seizure interval. This patient had too few seizures to meet the seizure clustering criteria. C) Lead seizures with a lead time of 1 day (gray) are plotted relative to the time of the seizure. The autocorrelation and variance did not tend to stay elevated after a lead seizure. The black lines shows the average with standard error bars. Red line shows a moving sum of seizures in a 2 hour window.

### Patient 6

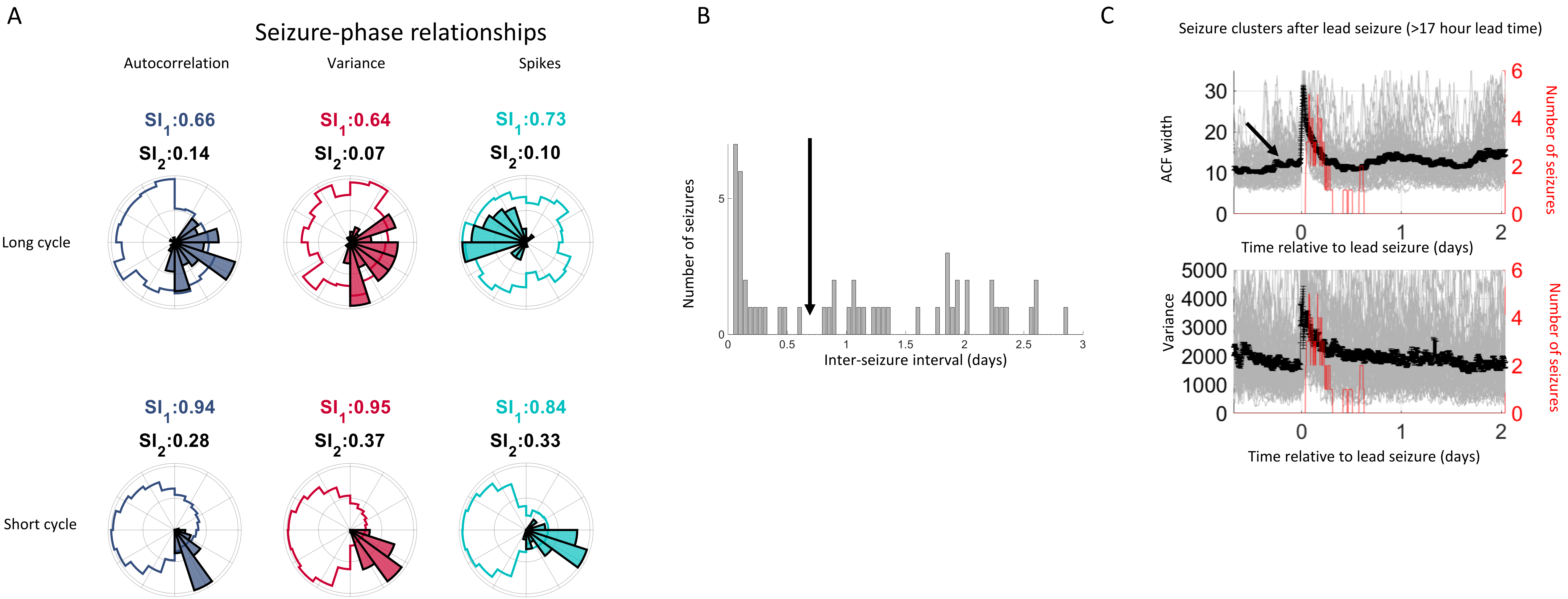


Supplementary Figure 6: Results summary for Patient 6. A) The Synchronization Indices ($SI_1$) for the autocorrelation (gray-blue), variance (red) and spikes (cyan) quantify the synchrony between seizures and the underlying signals. A high $SI_1$ corresponds to good alignment between seizures and the phase of the signal. The SI for the signals ($SI_2$) quantifies the phase uniformity of the signal. A low $SI_2$ demonstrates that the signal is periodic and that all phases are equally represented. B) The inter-seizure interval. This patient had peaks at approximately 1 and 2 days. We set the cut-off at 0.7 days (17 hours). C) Lead seizures with a lead time of 17 hours (gray) are plotted relative to the time of the seizure. The arrow denotes a visible increase in autocorrelation prior to the seizure that is overshadowed by the seizure event itself. After a lead seizure, the autocorrelation and variance signals decayed to baseline over a few hours. During this period, there was an increase susceptibility to seizures. The black lines shows the average with standard error bars. Red line shows a moving sum of seizures in a 2 hour window.

### Patient 7

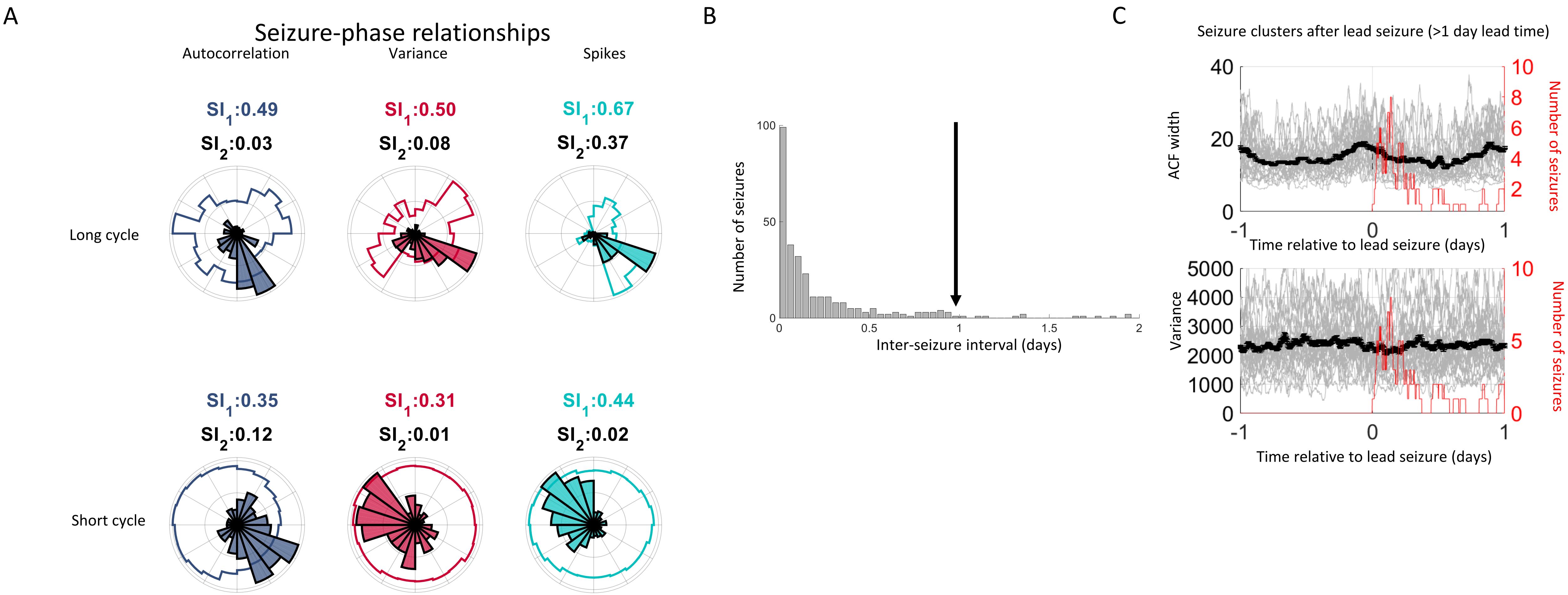


Supplementary Figure 7: Results summary for Patient 7. A) The Synchronization Indices ($SI_1$) for the autocorrelation (gray-blue), variance (red) and spikes (cyan) quantify the synchrony between seizures and the underlying signals. A high $SI_1$ corresponds to good alignment between seizures and the phase of the signal. The SI for the signals ($SI_2$) quantifies the phase uniformity of the signal. A low $SI_2$ demonstrates that the signal is periodic and that all phases are equally represented. B) The inter-seizure interval. This patient had an exponential decay in the histogram. We set the lead seizure cut-off at 1 day. C) Lead seizures with a lead time of 1 day (gray) are plotted relative to the time of the seizure. After a lead seizure, the autocorrelation quickly decayed. Despite this steady decrease, there was an increased seizure susceptibility. There was no obvious change in the signal variance. The black lines shows the average with standard error bars. Red line shows a moving sum of seizures in a 2 hour window.

### Patient 8

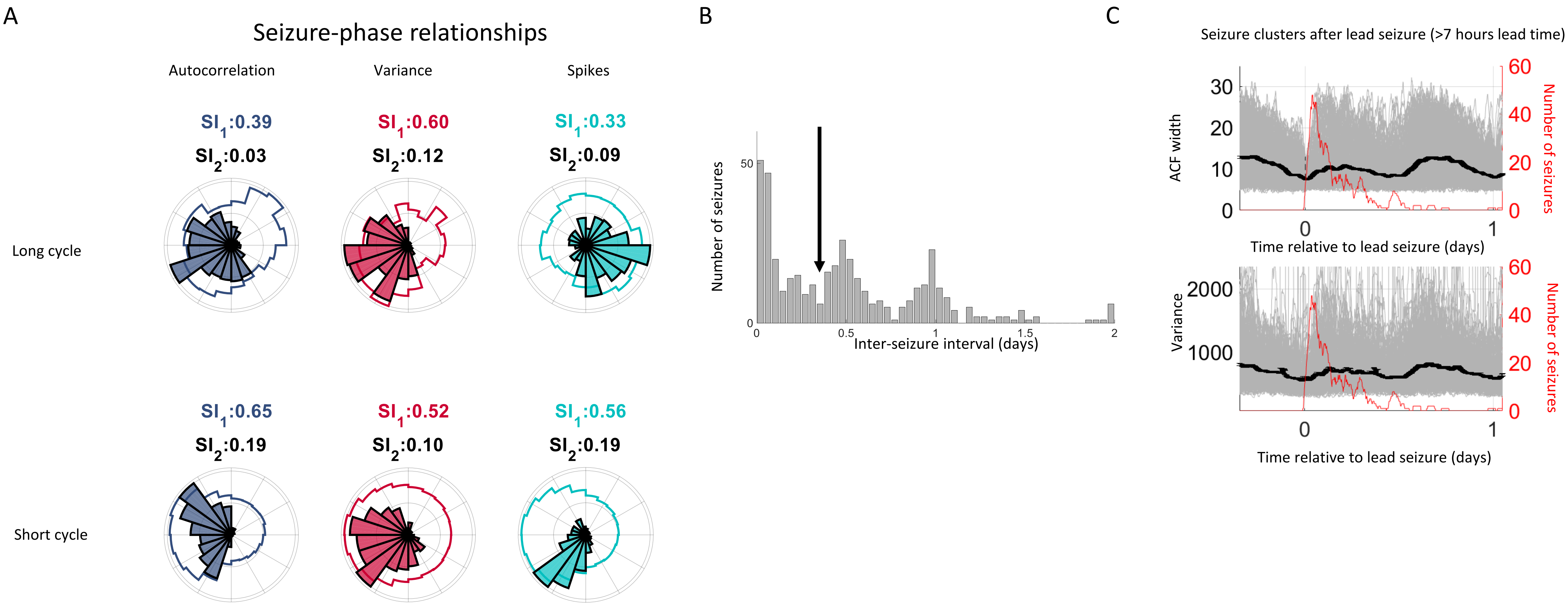


Supplementary Figure 8: Results summary for Patient 8. A) The Synchronization Indices ($SI_1$) for the autocorrelation (gray-blue), variance (red) and spikes (cyan) quantify the synchrony between seizures and the underlying signals. A high $SI_1$ corresponds to good alignment between seizures and the phase of the signal. The SI for the signals ($SI_2$) quantifies the phase uniformity of the signal. A low $SI_2$ demonstrates that the signal is periodic and that all phases are equally represented. B) The inter-seizure interval. This patient had multiple peaks separated by approximately 12 hours. We set the lead seizure cut-off to 0.3 days (7 hours). C) Lead seizures with a lead time of 7 hours (gray) are plotted relative to the time of the seizure. After a lead seizure, the autocorrelation and variance signals steadily increased. During this time there was an increased seizure susceptibility. The black lines shows the average with standard error bars. Red line shows a moving sum of seizures in a 2 hour window.

### Patient 9

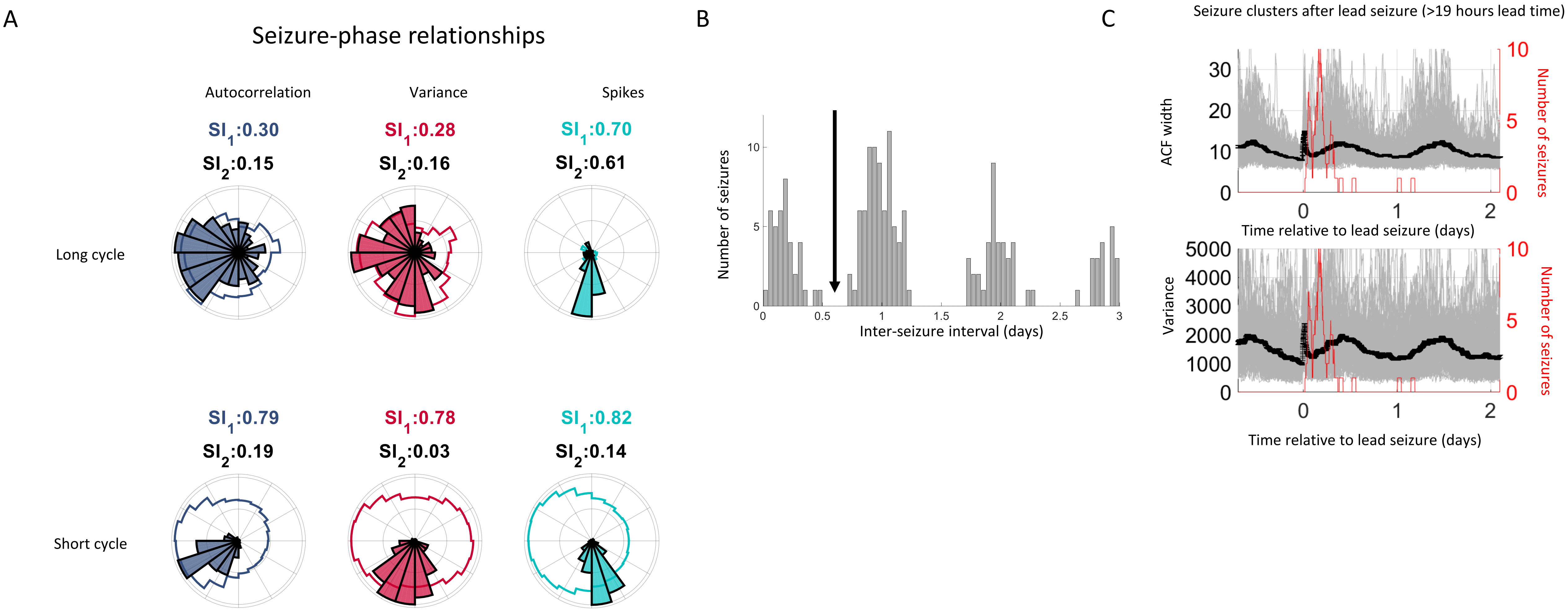


Supplementary Figure 9: Results summary for Patient 9. A) The Synchronization Indices ($SI_1$) for the autocorrelation (gray-blue), variance (red) and spikes (cyan) quantify the synchrony between seizures and the underlying signals. A high $SI_1$ corresponds to good alignment between seizures and the phase of the signal. The SI for the signals ($SI_2$) quantifies the phase uniformity of the signal. A low $SI_2$ demonstrates that the signal is periodic and that all phases are equally represented. B) The inter-seizure interval. This patient had multiple peaks separated by approximately 1 day. We set the lead seizure cut-off to 0.7 days (19 hours). C) Lead seizures with a lead time of 19 hours (gray) are plotted relative to the time of the seizure. After a lead seizure, the autocorrelation and variance signals steadily increased. During this time there was an increased seizure susceptibility. The black lines shows the average with standard error bars. Red line shows a moving sum of seizures in a 2 hour window.

### Patient 10

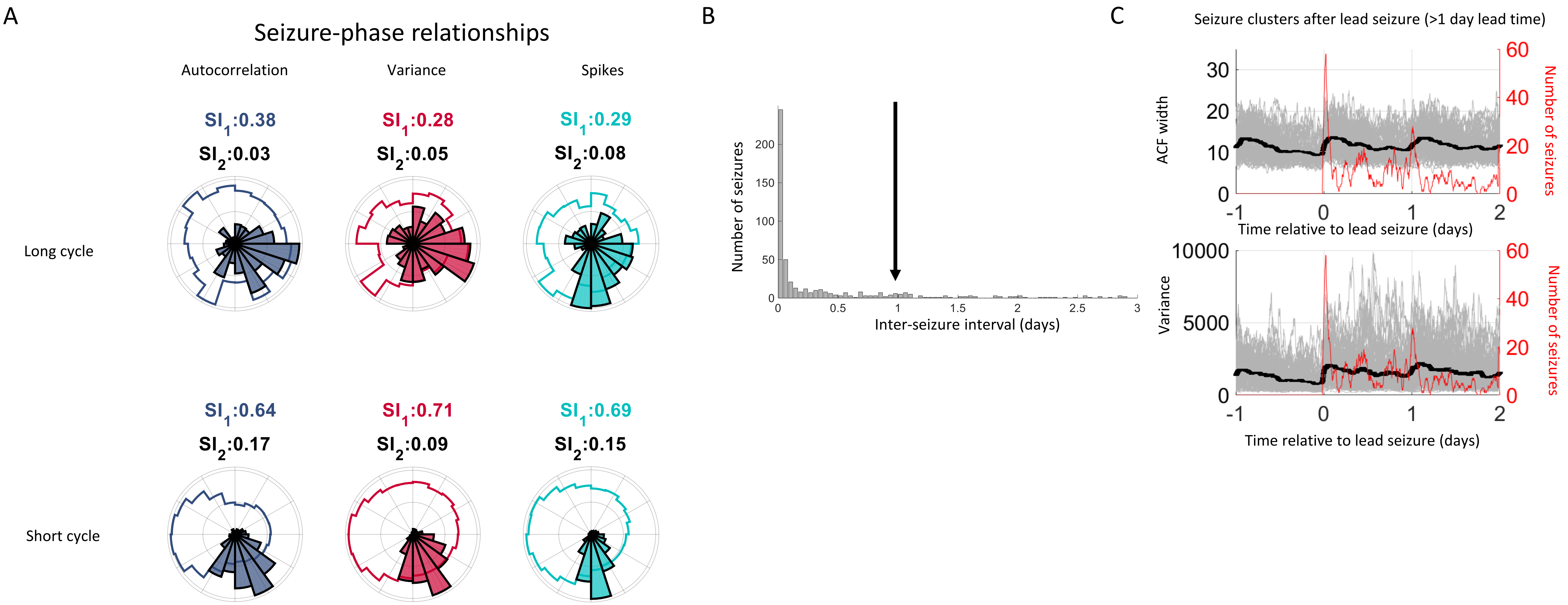


Supplementary Figure 10: Results summary for Patient 10. A) The Synchronization Indices ($SI_1$) for the autocorrelation (gray-blue), variance (red) and spikes (cyan) quantify the synchrony between seizures and the underlying signals. A high $SI_1$ corresponds to good alignment between seizures and the phase of the signal. The SI for the signals ($SI_2$) quantifies the phase uniformity of the signal. A low $SI_2$ demonstrates that the signal is periodic and that all phases are equally represented. B) The inter-seizure interval. This patient had an exponential decay in the histogram. We set the lead seizure cut-off at 1 day. C) Lead seizures with a lead time of 1 day (gray) are plotted relative to the time of the seizure. After a lead seizure, the autocorrelation and variance signals steadily increased. During this time there was an increased seizure susceptibility. The black lines shows the average with standard error bars. Red line shows a moving sum of seizures in a 2 hour window.

### Patient 11

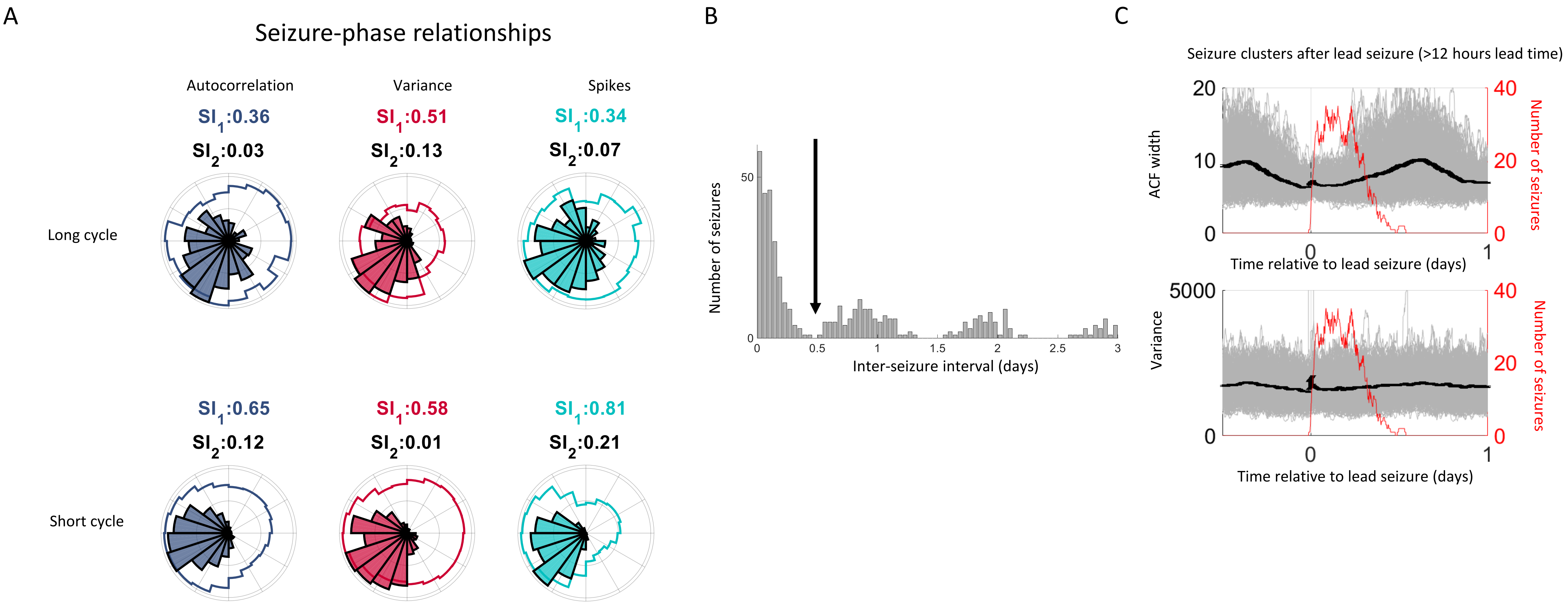


Supplementary Figure 11: Results summary for Patient 11. A) The Synchronization Indices ($SI_1$) for the autocorrelation (gray-blue), variance (red) and spikes (cyan) quantify the synchrony between seizures and the underlying signals. A high $SI_1$ corresponds to good alignment between seizures and the phase of the signal. The SI for the signals ($SI_2$) quantifies the phase uniformity of the signal. A low $SI_2$ demonstrates that the signal is periodic and that all phases are equally represented. B) The inter-seizure interval. This patient had multiple peaks separated by approximately 1 day. We set the lead seizure cut-off to 0.5 days (12 hours). C) Lead seizures with a lead time of 12 hours (gray) are plotted relative to the time of the seizure. After a lead seizure, the autocorrelation signal steadily increased. During this time there was an increased seizure susceptibility. There was not any obvious change in the signal variance. The black lines shows the average with standard error bars. Red line shows a moving sum of seizures in a 2 hour window.

### Patient 12

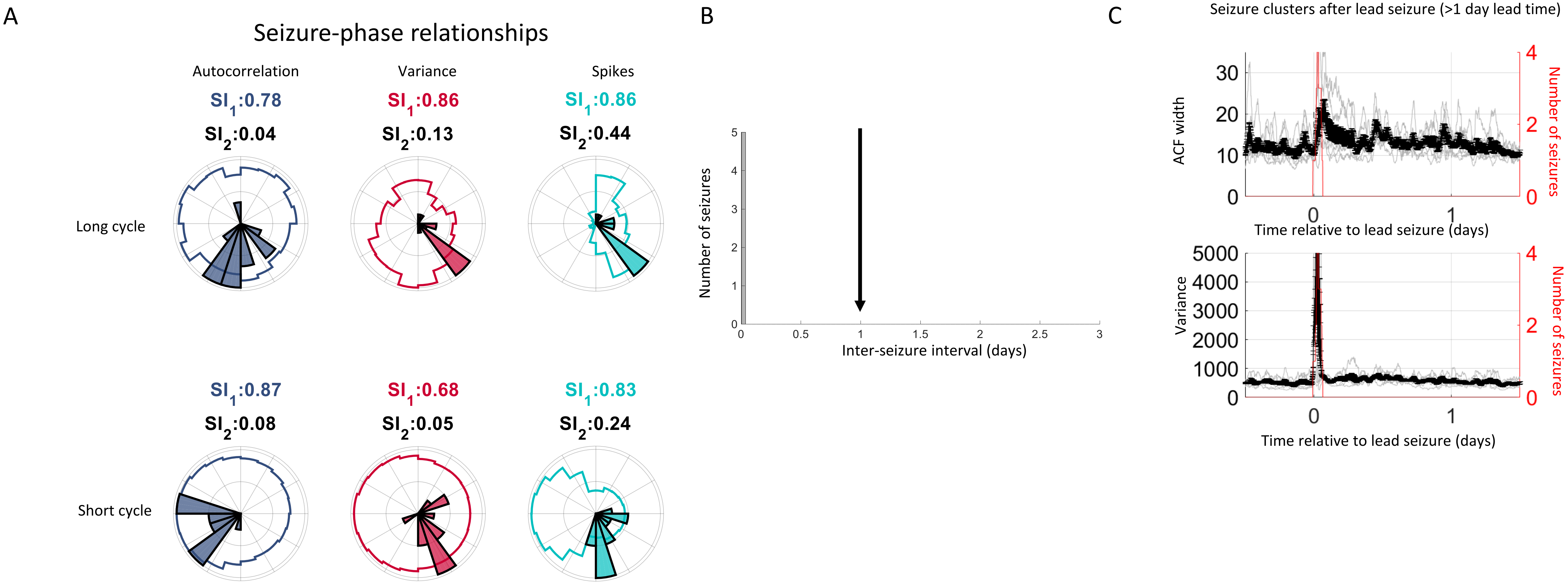


Supplementary Figure 12: Results summary for Patient 12. A) The Synchronization Indices ($SI_1$) for the autocorrelation (gray-blue), variance (red) and spikes (cyan) quantify the synchrony between seizures and the underlying signals. A high $SI_1$ corresponds to good alignment between seizures and the phase of the signal. The SI for the signals ($SI_2$) quantifies the phase uniformity of the signal. A low $SI_2$ demonstrates that the signal is periodic and that all phases are equally represented. B) The inter-seizure interval showed that a very high proportion of seizures occurred within 1 hour of a previous seizure. We set the lead seizure cut-off at 1 day. C) Lead seizures (gray) are plotted relative to the time of the seizure. After a lead seizure, the autocorrelation and variance signals decayed to baseline over a few hours. During this period, there was an increase susceptibility to seizures. The black lines shows the average with standard error bars. Red line shows a moving sum of seizures in a 2 hour window.

### Patient 13

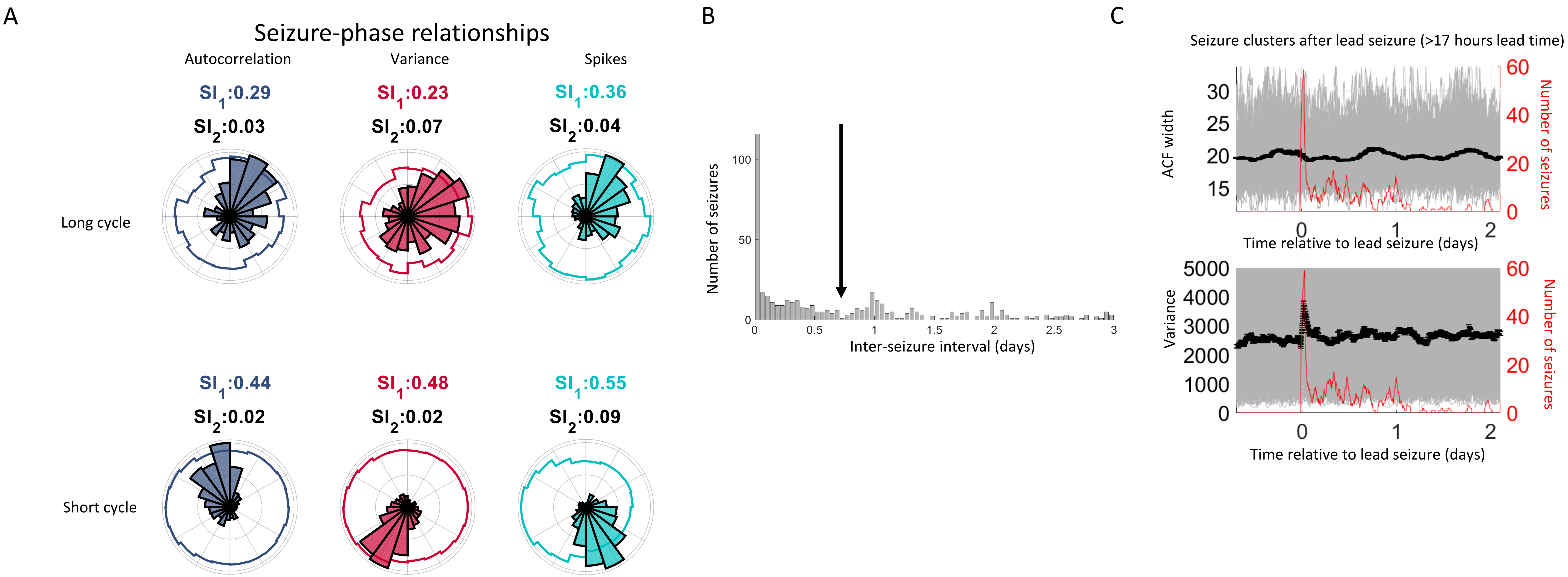


Supplementary Figure 13: Results summary for Patient 13. A) The Synchronization Indices ($SI_1$) for the autocorrelation (gray-blue), variance (red) and spikes (cyan) quantify the synchrony between seizures and the underlying signals. A high $SI_1$ corresponds to good alignment between seizures and the phase of the signal. The SI for the signals ($SI_2$) quantifies the phase uniformity of the signal. A low $SI_2$ demonstrates that the signal is periodic and that all phases are equally represented. B) The inter-seizure interval. This patient had peaks at approximately 1 and 2 days. We set the cut-off at 0.7 days (17 hours). C) Lead seizures with a lead time of 17 hours (gray) are plotted relative to the time of the seizure. After a lead seizure, the variance signals decayed to baseline over a few hours. During this period, there was an increase susceptibility to seizures. There was no obvious change in the autocorrelation signal. The black lines shows the average with standard error bars. Red line shows a moving sum of seizures in a 2 hour window.

### Patient 14

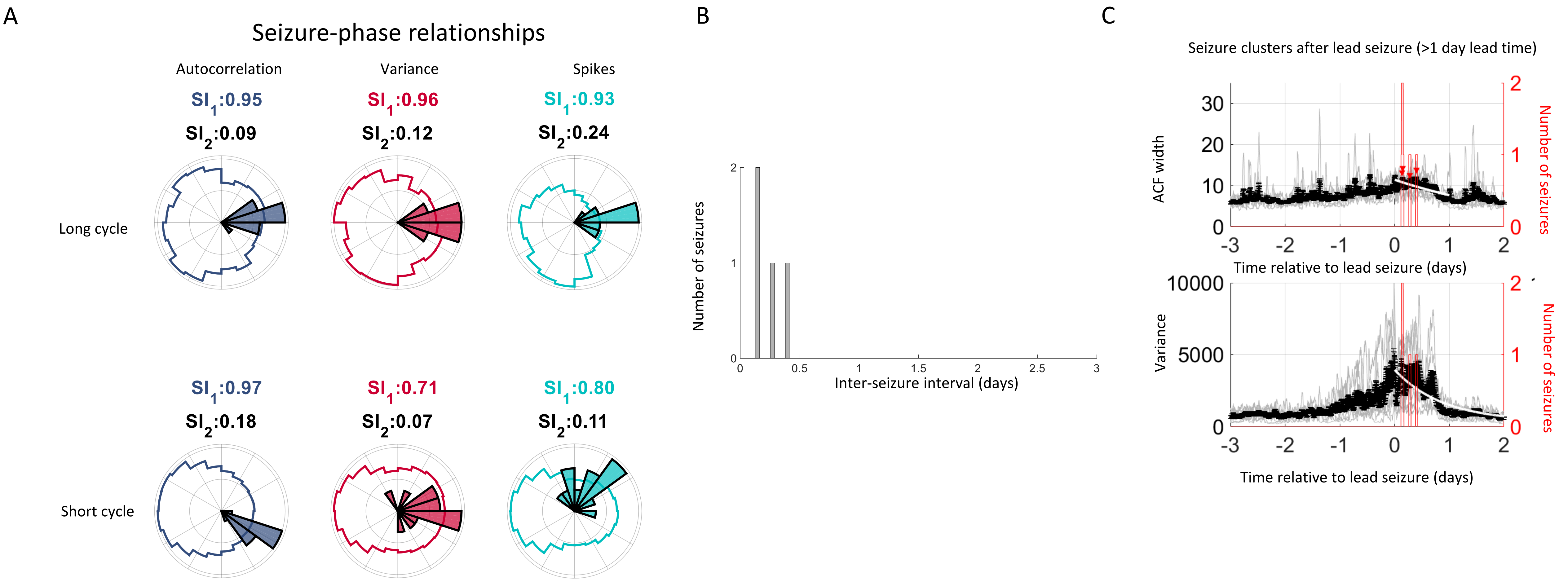


Supplementary Figure 14: Results summary for Patient 14. A) The Synchronization Indices ($SI_1$) for the autocorrelation (gray-blue), variance (red) and spikes (cyan) quantify the synchrony between seizures and the underlying signals. A high $SI_1$ corresponds to good alignment between seizures and the phase of the signal. The SI for the signals ($SI_2$) quantifies the phase uniformity of the signal. A low $SI_2$ demonstrates that the signal is periodic and that all phases are equally represented. B) The inter-seizure interval. This patient had too few seizures to meet the seizure clustering criteria. C) Lead seizures with a lead time of 1 day (gray) are plotted relative to the time of the seizure. There was an obvious increase in autocorrelation and variance that occurred over 1-2 days prior to a seizure. After a lead seizure, the two signals decayed to baseline over approximately 1 day. During this period, there was an increase susceptibility to seizures. The black lines shows the average with standard error bars. Red line shows a moving sum of seizures in a 2 hour window.

### Patient 15

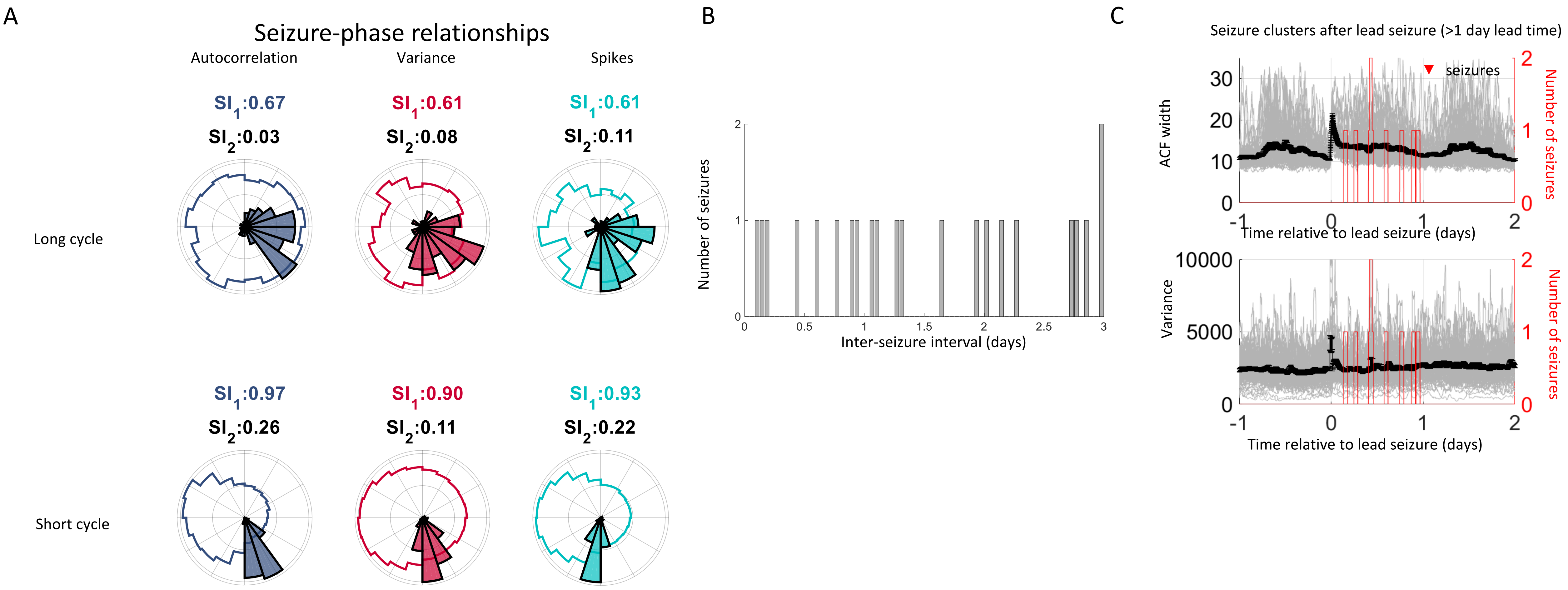


Supplementary Figure 15: Results summary for Patient 15. A) The Synchronization Indices ($SI_1$) for the autocorrelation (gray-blue), variance (red) and spikes (cyan) quantify the synchrony between seizures and the underlying signals. A high $SI_1$ corresponds to good alignment between seizures and the phase of the signal. The SI for the signals ($SI_2$) quantifies the phase uniformity of the signal. A low $SI_2$ demonstrates that the signal is periodic and that all phases are equally represented. B) The inter-seizure interval. This patient had too few seizures to meet the seizure clustering criteria. C) Lead seizures with a lead time of 1 day (gray) are plotted relative to the time of the seizure. After a lead seizure, the autocorrelation signal decayed to baseline over a few hours. Seizures tended to occur randomly irrespective of the autocorrelation and variance signals. The black lines shows the average with standard error bars. Red line shows a moving sum of seizures in a 2 hour window.

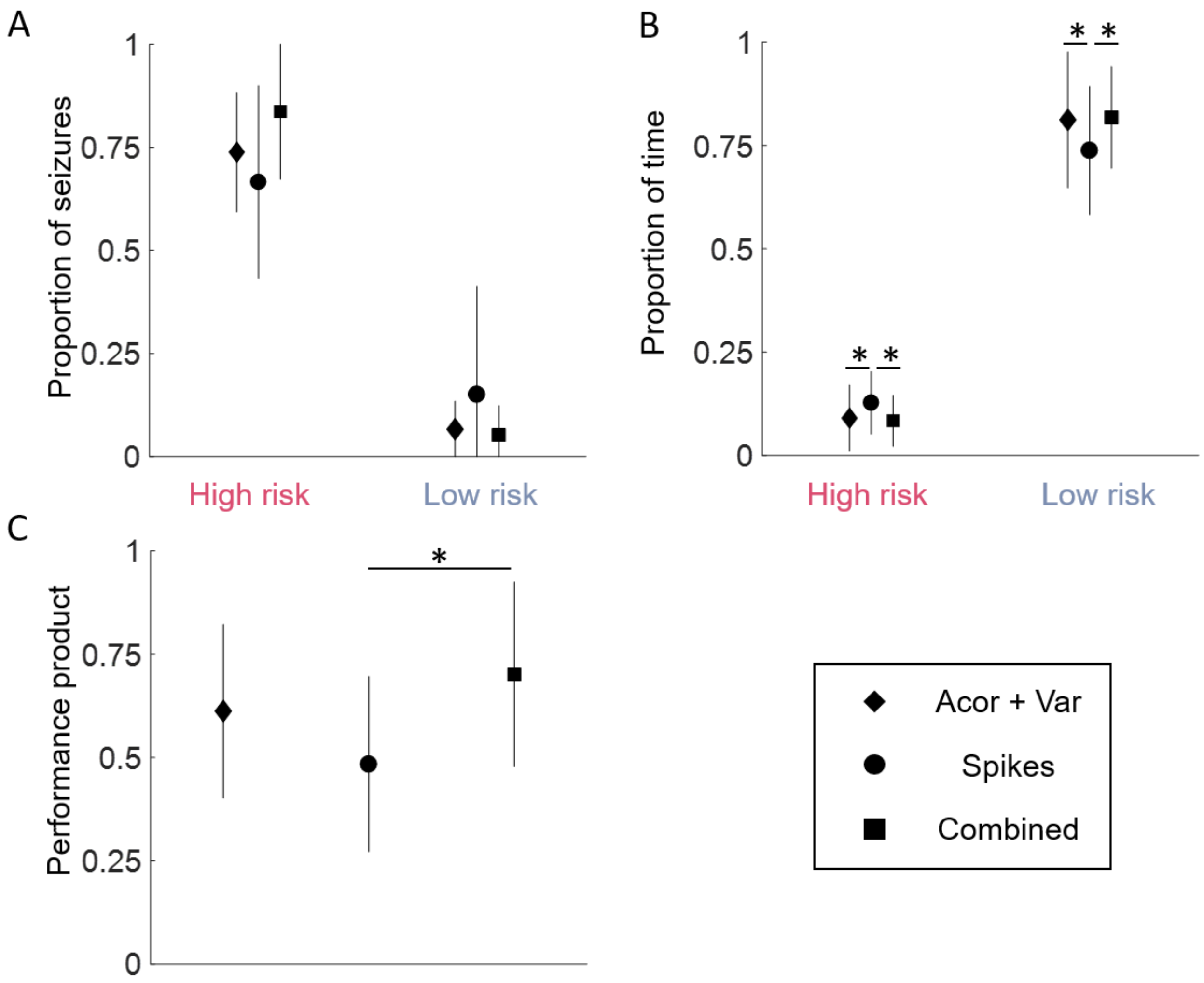


Supplementary Figure 16: Forecasting performance is compared for critical slowing (autocorrelation and variance), spikes and the combination of the three measures. A) there were no significant differences in the number of seizures correctly classified as high risk, or the number of seizures occurring during low risk. B) There was a significantly higher proportion of time spent in the high risk state, and significantly lower amount of time in the low risk state using the spike rate model than the other two models. C) Overall, the combined model performed significantly better than the spike model. Significance is measured to $p < 0.01$.

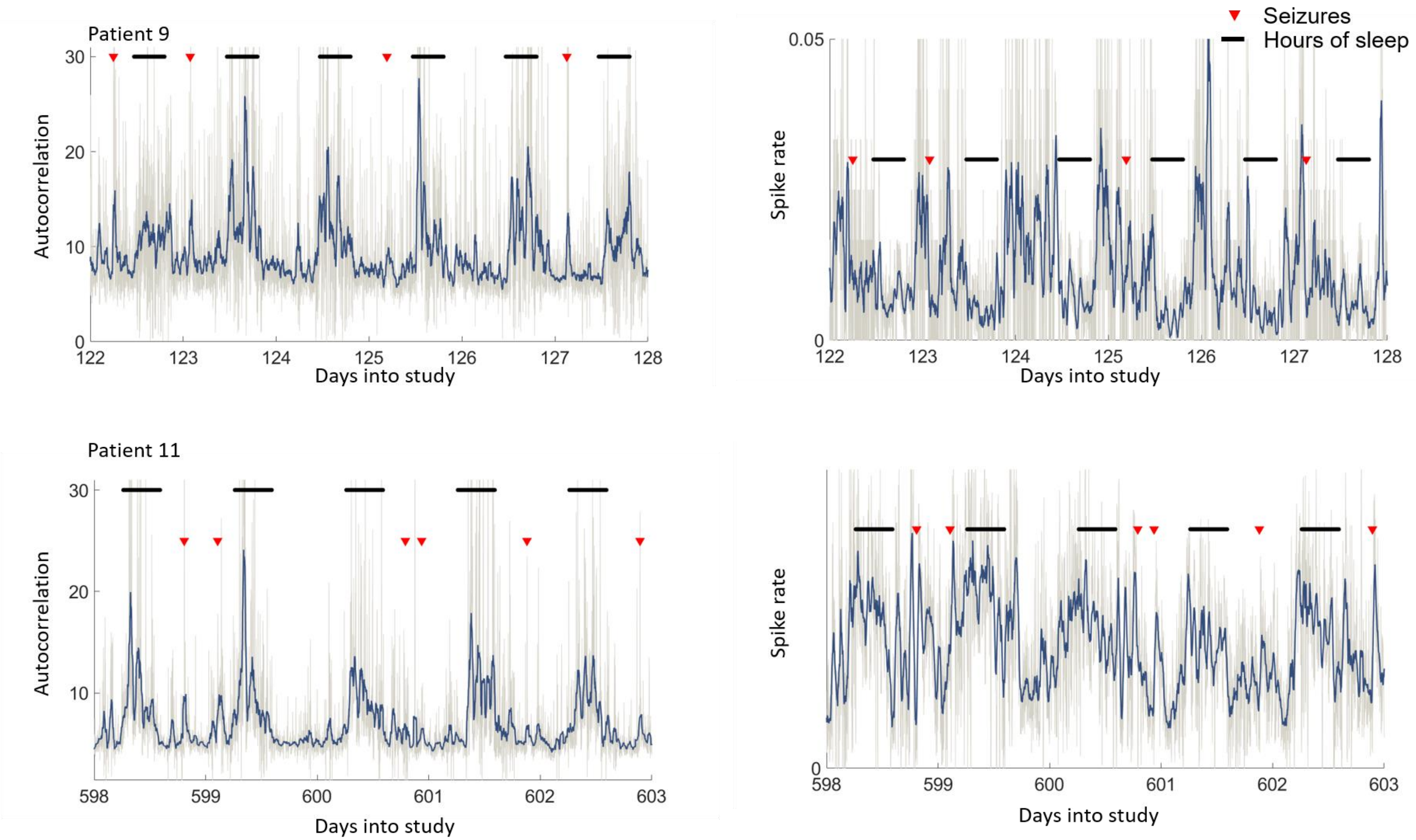


Supplementary Figure 17: The autocorrelation signal is a marker of a state change. However, there are possibly many state changes that occur in the brain. For example, the sleep-wake is characterised by a large increase in autocorrelation. These examples for patient 9 and 11 demonstrate how the autocorrelation increased by a large amount every sleep-wake cycle (black bars). Seizures, on the other hand, were generally characterised by small increases in autocorrelation. In these two patients, seizures were preceded by large increases in spike rate. Hours of sleep from 10 pm to 6 am.

Table 2: Summary of forecasting results

| Patient | Method | Seizures in Low | Seizures in High | Time in Low | Time in High |
|---|---|---|---|---|---|
| 1 | M1 | 3 | 91 | 95 | 3 |
| | M2 | 9 | 88 | 100 | 8 |
| | OT† training | | 75 | 27 | 33 |
| | OT advisory | | 77 | 7 | 27 |
| 2 | M1 | 9 | 78 | 93 | 3 |
| | M2 | 9 | 88 | 100 | $2\times10^{-3}$ |
| | OT training | | 75 | 58 | 21 |
| | OT advisory | | 100 | 56 | 31 |
| 4 | M1 | 9 | 86 | 99 | 0.6 |
| | M2 | 14 | 73 | 100 | $2\times10^{-3}$ |
| 6 | M1 | 0 | 94 | 99 | 0.6 |
| | M2 | 27 | 66 | 95 | 3 |
| 7 | M1 | 13 | 69 | 79 | 10 |
| | M2 | 22 | 64 | 66 | 23 |
| 8 | M1 | 15 | 61 | 72 | 13 |
| | M2 | 12 | 72 | 83 | 11 |
| | OT training | | 63 | | 40 |
| | OT advisory | | 62 | | 28 |
| 9 | M1 | 2 | 88 | 69 | 15 |
| | M2 | 7 | 85 | 81 | 16 |
| | OT training | | 59 | 19 | 36 |
| | OT advisory | | 17 | 48 | 11 |
| 10 | M1 | 7 | 80 | 77 | 11 |
| | M2 | 11 | 79 | 66 | 24 |
| | OT training | | 75 | | 31 |
| | OT advisory | | 51 | | 17 |
| 11 | M1 | 6 | 76 | 76 | 12 |
| | M2 | 6 | 86 | 81 | 16 |
| | OT training | | 65 | 20 | 30 |
| | OT advisory | | 39 | 26 | 15 |
| 12 | M1 | 0 | 71 | 100 | $2\times10^{-4}$ |
| | M2 | 15 | 85 | 100 | $2\times10^{-4}$ |
| 13 | M1 | 10 | 70 | 67 | 14 |
| | M2 | 11 | 69 | 79 | 10 |
| | OT training | | 62 | | 35 |
| | OT advisory | | 50 | | 28 |
| 14 | M1 | 0 | 73 | 100 | $5\times10^{-4}$ |
| | M2 | 8 | 67 | 100 | $2\times10^{-3}$ |
| 15 | M1 | 0 | 97 | 99 | 0.8 |
| | M2 | 6 | 87 | 89 | 7 |
| | OT training | | 100 | | 18 |
| | OT advisory | | 71 | | 41 |
| Total‡ | M1 | 5 ± 7 | 84 ± 16 | 82 ± 12 | 8 ± 6 |
| | M2 | 13 ± 6 | 77 ± 8 | 87 ± 12 | 9 ± 8 |
| | OT training | | 72 ± 13 | 31 ± 18 | 31 ± 8 |
| | OT advisory | | 58 ± 25 | 34 ± 22 | 25 ± 10 |

All values in the table represent percentages rounded to the closest integer.

† OT – original trial. Where available, sensitivity (seizures in high), time in high and time in low values from the original trial in the training and advisory phases are shown [21].

‡ Values represent mean ± 1 standard deviation.

Supplementary Figure 18: Summary of forecasting results for Method M1 and M2. Where available, these results are compared to the original trial [21], which included a training and advisory phase.

#### Comparison of NeuroVista data studies – Sensitivity (S)/Time in high (TiH)

| Patient | Original [21] | | Deep CNN [31] | | CW [32] | | Logistic [32] | | CW + Logistic [32] | | Kaggle [33] | | CW + Kaggle [33] | | Critical slowing | |
|---|---|---|---|---|---|---|---|---|---|---|---|---|---|---|---|---|
| | S | TiH | S | TiH | S | TiH | S | TiH | S | TiH | S | TiH | S | TiH | S | TiH |
| 1 | 77 | 27 | 65 | 21 | 34 | 27 | 54 | 27 | 61 | 27 | | | | | 83 | 8 |
| 2 | 100 | 31 | 74 | 11 | | | | | | | | | | | 87 | 0.02 |
| 3 | 45 | 29 | 71 | 53 | 36 | 29 | 53 | 29 | 55 | 29 | 66 | 29 | 60 | 29 | | |
| 4 | | | | | | | | | | | | | | | 72 | 0.02 |
| 6 | | | | | 52 | | 61 | | 65 | | | | | | 66 | 3 |
| 7 | | | | | | | | | | | | | | | 76 | 21 |
| 8 | 62 | 28 | 77 | 32 | 58 | 28 | 71 | 28 | 76 | 28 | | | | | 64 | 23 |
| 9 | 17 | 11 | 83 | 43 | 28 | 11 | 29 | 11 | 45 | 11 | 39 | 11 | 52 | 11 | 85 | 16 |
| 10 | 51 | 17 | 68 | 32 | 36 | 17 | 38 | 17 | 52 | 17 | 48 | 17 | 53 | 17 | 78 | 24 |
| 11 | 39 | 15 | 78 | 18 | 43 | 15 | 57 | 15 | 58 | 15 | | | | | 86 | 16 |
| 12 | | | | | | | | | | | | | | | 85 | $2\times10^{-4}$ |
| 13 | 50 | 28 | 70 | 21 | 61 | 28 | 78 | 28 | 76 | 28 | | | | | 64 | 14 |
| 14 | 100 | 3 | 42 | 2 | | | | | | | | | | | 69 | $6\times10^{-4}$ |
| 15 | 71 | 21 | 59 | 37 | 71 | 21 | 51 | 21 | 60 | 21 | | | | | 87 | 0.07 |
| Average | 61 (52) | 21 (22) | 69 (71) | 27 (32) | (47) | (21) | (57) | (21) | (61) | (21) | | | | | 78 (77) | 9 (11) |

- Best method in green
- Averages in parentheses ignore patients 2 and 14

Supplementary Figure 19: Comparison of the prospective forecasting results (Method M2) and previously developed prospective forecasting methods using the same dataset.
